# Comparison between the activities of canonical Wnt ligands in human pluripotent stem cell differentiation

**DOI:** 10.64898/2025.12.18.695264

**Authors:** Eleni Anastasia Rizou, Aryeh Warmflash

## Abstract

WNT ligands are key regulators of mammalian gastrulation and anterior-posterior (A–P) patterning. The activities of different Wnt ligands that signal through the canonical pathway are often assumed to be similar due to their high structural homology. To explore potential differences, we generated human pluripotent stem cell (hPSC) lines with inducible expression of WNT3, WNT3A, WNT6, or WNT8A, enabling direct comparisons of ligand activity in multiple differentiation contexts. We find that only WNT3 and WNT3A robustly induce mesodermal markers, while WNT6 and WNT8A do not. In the context of ectodermal differentiation, WNT3, WNT3A, and WNT8 activate a similar set of targets but WNT3 and 3A do so more strongly than WNT8A. WNT6 does not induce this set of targets, however, all four ligands, including WNT6, downregulate a similar set of targets. We also observed ligand-specific differences in spatial signaling range and the ability to induce self-organized patterns of ectodermal fates including neural crest-like domains. These results highlight how signaling strength, spatial range, and diverse target genes of Wnt ligands contribute to distinct developmental outcomes in early human stem cell differentiation.

## Introduction

WNT ligands are essential regulators of embryonic development, governing diverse processes including axis formation, cell fate specification, and tissue patterning. In vertebrates, the 19 conserved WNT ligands are broadly categorized into canonical and non-canonical classes based on their ability to activate *β* -catenin–dependent transcriptional programs. Canonical WNT signaling stabilizes *β*-catenin and leads to activation of TCF/LEF transcription factors, while non-canonical pathways engage alternative intracellular mechanisms such as the planar cell polarity and calcium signaling pathways (Logan and Nusse 2004; Niehrs 2012; Boutros and Niehrs 2016; Nusse and Clevers 2017). However, these functional categories are not absolute: several WNT ligands can activate both pathways in a context-dependent manner (Mikels and Nusse 2006; Clevers 2006; Amerongen and Nusse 2009).

In early mammalian development, WNT signaling plays indispensable roles during gastrulation and neurulation. Mouse knockout studies of Wnt3, *β*-catenin, or Porcn (an enzyme required for WNT secretion) result in early embryonic arrest at the onset of gastrulation, demonstrating a requirement for canonical WNT signaling in primitive streak formation (Haegel et al. 1995; Liu et al. 1999; Barrott et al. 2011; Biechele et al. 2013). WNT3 and WNT3A are central to mesoderm induction and posterior body formation, while WNT8A and WNT6 have been associated with patterning roles during later stages of neurulation and neural crest emergence in various model systems (Christian et al. 1991; Takada et al. 1994; García-Castro et al. 2002; Cunningham et al. 2015; Wylie et al. 2014; Wei et al. 2020).

Despite extensive functional studies on individual ligands, the degree to which different WNTs are functionally interchangeable remains unclear. Their evolutionary conservation and shared classification as canonical ligands have led to assumptions of redundancy, but emerging evidence suggests distinct ligand–receptor interactions, signaling kinetics, and diffusion ranges (Dijksterhuis et al. 2015; Voloshanenko et al. 2017; Tao et al. 2019). Comparative studies directly testing the specificity of WNT ligands in a shared cellular context remain rare, due in part to technical challenges in standardizing ligand delivery and readouts (Tanaka et al. 2011; Chhabra et al. 2019).

To understand the similarities and differences between the activity induced by different Wnt ligands, we systematically examined the outcome of signaling through four different ligands — WNT3, WNT3A, WNT6, and WNT8A — during the differentiation of hPSCs to primitive streak or ectodermal fates. Using hPSCs engineered to inducibly express each ligand, we compared their abilities to activate canonical signaling and induce posterior and mesodermal cell fates. In addition, we explored the signaling range of each ligand, and how engineered spatial gradients create patterns of cell fates during ectodermal differentiation. These analyses provide a side-by-side comparison of WNT ligand function and show that different Wnt ligands which signal through the canonical pathway are not functionally equivalent but differ in strength, targets, and range of action.

## Results

### WNT3, WNT3A and WNT8A induce TCF/LEF signaling with different strengths

We chose to study four different Wnt ligands, WNT3, WNT3A, WNT8A, and WNT6, due to their roles in early development described above, and their expression in 2D-gastruloids, micropatterned hPSC colonies treated with BMP4, which differentiate in spatial patterns consisting of amnion and all three embryonic germ layers (Warmflash et al. 2014; Chhabra et al. 2019; Tyser et al. 2020; Ferretti and Hadjantonakis 2019; Minn et al. 2021) (1A). These ligands show significant similarity in their sequence and structure which led us to examine the degree of similarity in their signaling activity. We generated four human embryonic stem cell lines, derived from the ESI017 line, each containing a cassette for overexpression of a WNT ligand, and a nuclear RFP separated by a self-cleaving t2a peptide and under the control of a doxycycline (dox)-inducible promoter (Figure 1B).

**Figure 1:**
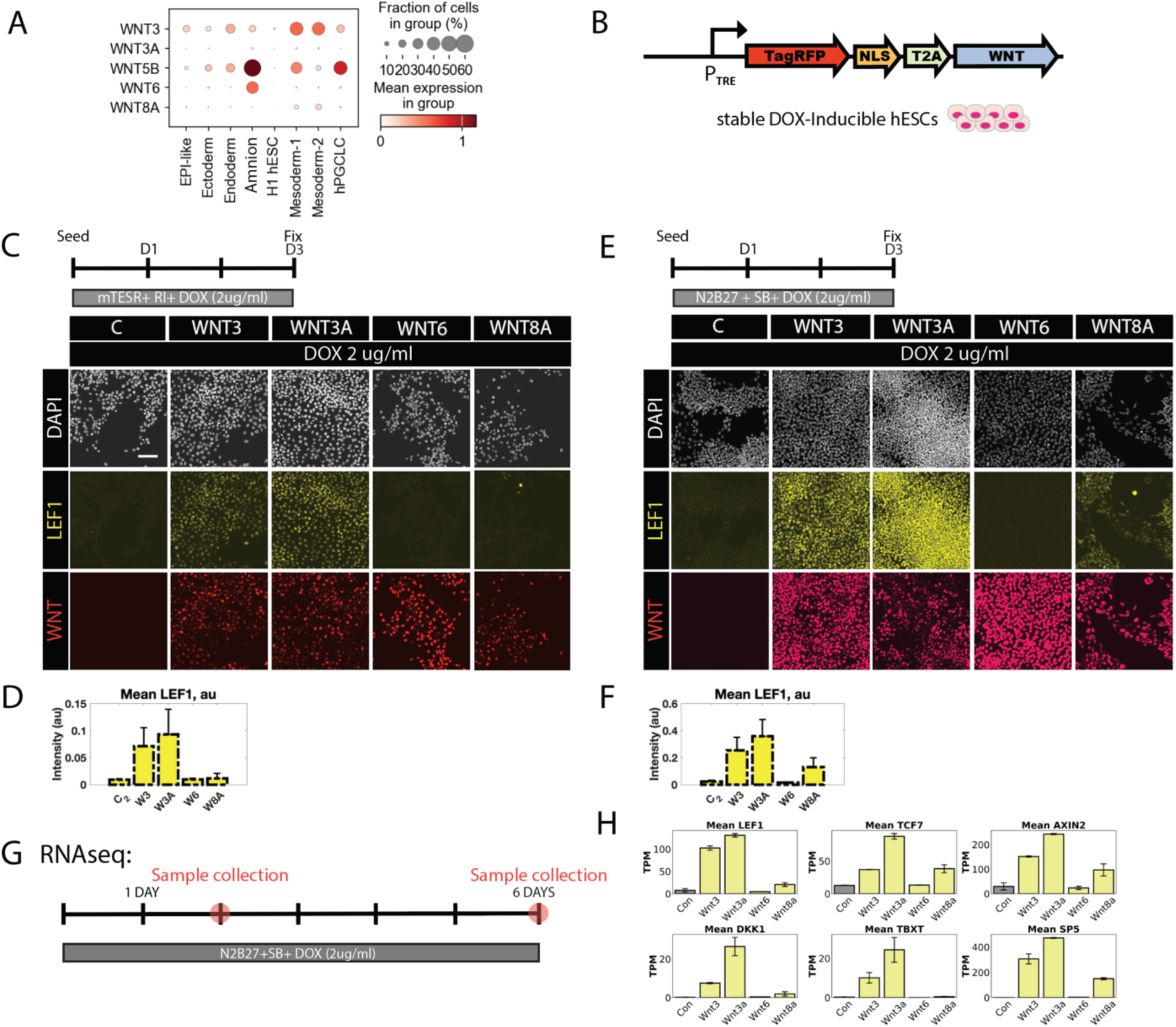
WNT3, WNT3A, and WNT8A induce canonical WNT targets with different strengths. (A). WNT ligand expression in micropatterned 2D human gastruloids. Data from (Minn et al. 2021). (B). Schematic of the constructs used to make the WNT-expressing stable doxycycline-induced cell lines. (C,E). The indicated cells lines were treated with doxycycline for 3 days either in mTeSR1 (C) or N2B27 (E), fixed and stained by immunofluorescence for LEF1. (D,F). Quantification of normalized fluorescence levels for the experiments shown in C (D) or E (F). (G). Schematic of RNA sequencing experiments. (H). Quantification of the indicated WNT targets from RNA sequencing data.

We sought to determine the degree to which different WNT ligands activate canonical WNT signaling when overexpressed. We monitored the expression of LEF1 and TBXT, two direct targets of canonical WNT signaling, following induction of WNT ligand expression with doxycycline (Figure S1, A,B). We first performed this experiment in the pluripotency media mTeSR1. Under these conditions, activation of canonical Wnt signaling leads to differentiation to primitive streak and subsequently to mesodermal fates. We found that both WNT3 and WNT3A overexpression activated LEF1 and TBXT along with other primitive streak and mesodermal fate markers, including CDX2 and MIXL1. In contrast, WNT8A and WNT6 did not induce LEF1 or TBXT (Figure 1C,Figure S1, A, B). Early endodermal markers, such as SOX17 and FOXA2, were expressed in only a few cells following WNT3 or WNT3A overexpression (Figure S2).

We also studied the action of Wnt overexpression in ectodermal differentiation. In this assay, cells are grown in N2B27 media supplemented with the TGF*β*/Activin/Nodal inhibitor SB431542 (Vallier et al. 2005). In this context, Wnt promotes the posteriorization of the neural ectoderm marked by downregulation of OTX2 and upregulation of HOX genes (Lippmann et al. 2015). Consistent with cells undergoing neural differentiation, all conditions, including controls, downregulated OCT4 and NANOG, while maintaining SOX2 at varying levels which depended on the WNT ligand(Figure S3, F-J). Under these conditions, WNT3 and WNT3A activated both LEF1 and TBXT. WNT8A activated LEF1 more weakly and we did not detect activation of TBXT. WNT6 overexpression did not lead to activation of either target (1D). Thus, WNT8A activated LEF1 in the context of ectodermal differentiation but not in pluripotency (Figure S1C, D).

To comprehensively examine gene expression and cell fates during ectodermal differentiation, we performed bulk RNA sequencing on all WNT-overexpressing cell lines at days 2 and 6 of doxycycline treatment (Figure 1E, F). Consistent with the results above, WNT3 and WNT3A induced high and comparable LEF1 expression, WNT8A induced low levels of LEF1 expression, and WNT6 induced none. This trend was mirrored across multiple canonical WNT target genes, such as TCF7, DKK1, SP5, and AXIN2, with WNT3A showing slightly stronger induction than WNT3 for most targets except LEF1 (Figure 1G). Mesodermal markers were highly expressed in WNT3 and WNT3A overexpressing cells on day 2 and minimally expressed by day 6 (Figure S1G). The posterior marker CDX2 was highly expressed by day 2 in WNT3 and WNT3A cells (Figure S1F). In agreement with the immunostaining results above, pluripotency markers were downregulated across all conditions, with the exception of SOX2, which persisted but was lower in WNT6 overexpressing cells (Figure S3G, K, L).

### WNT ligands posteriorize the neural ectoderm to different degrees

WNT ligands act as posteriorizing agents within the neuroectoderm and are critical for the development of cells at all axial positions from midbrain to spinal cord (Yamaguchi 2001; Green et al. 2015; Niehrs 2022). We determined the extent to which overexpression of each WNT ligand posteriorized cells during neural differentiation. Immunostaining of cells after 6 days of differentiation with doxycycline revealed that all four WNT ligands tested effectively suppressed the anterior marker OTX2 (Figure 2B,C). The same trend was observed with the anterior marker PAX6 (Figure S4 A,B). WNT3, WNT3A, and WNT8A, but not WNT6, induced the posterior marker HOXB1, with WNT3 and WNT3A showing the most robust activation.

**Figure 2:**
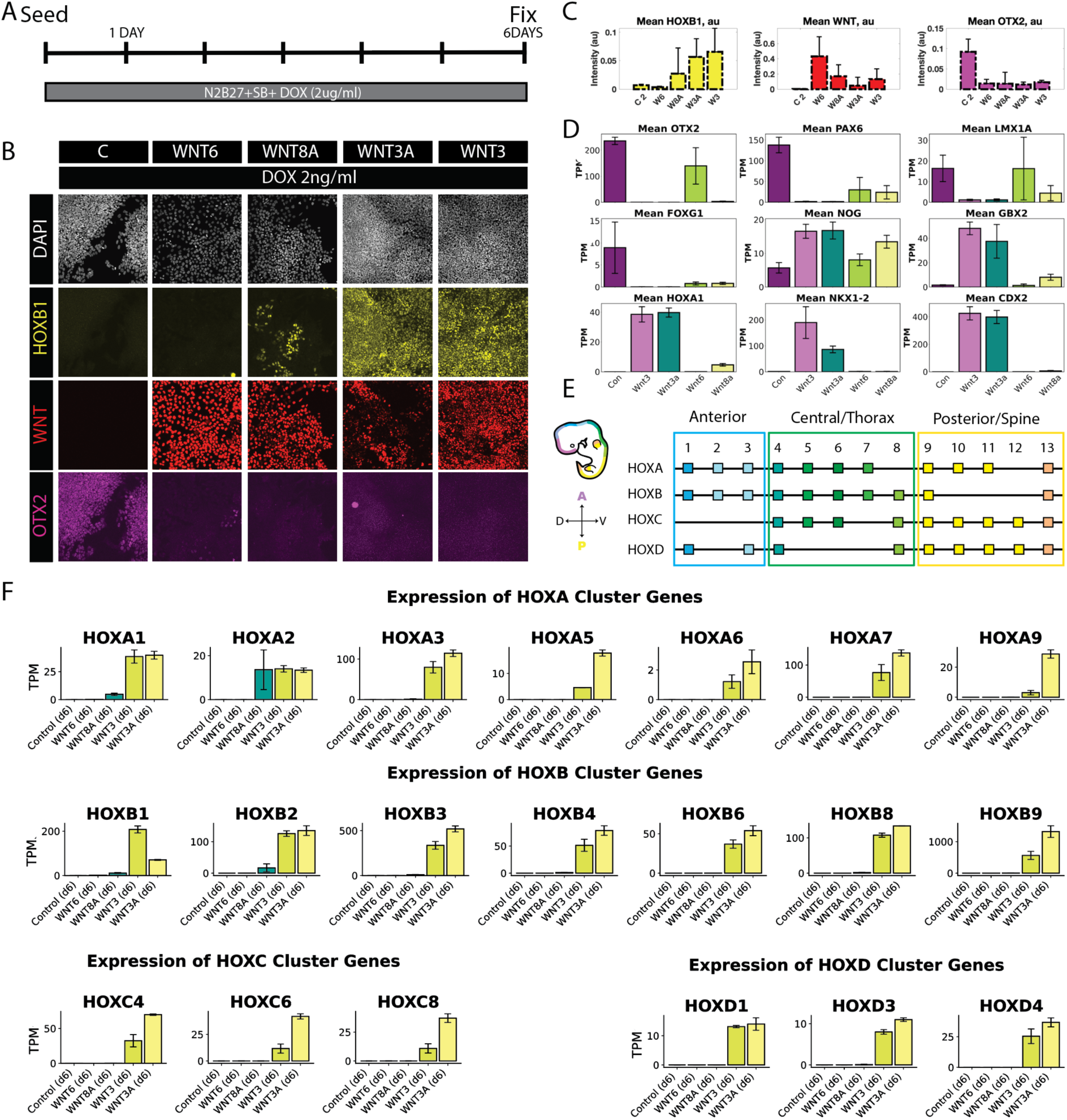
All WNT ligands suppress anterior fates but induce posterior fates with different strengths. (A). Schematic of differentiation protocol. (B). Immunostaining for OTX2 and HOXB1 after 6 days of neural differentiation with doxycycline (C). Quantification of the results in B. Error bars represent standard deviation across images (n = 6 images/condition). (D). Quantification of expression of the indicated AP markers after 6 days of differentiation from RNA-sequencing data. (E). Schematic showing the anteroposterior distribution of human HOX genes. (F). Quantification of HOX gene expression from RNA sequencing data.

We analyzed additional markers of AP position in the RNA sequencing data. All ligands suppressed the expression of anterior markers such as PAX6 and FOXG1. WNT3 and WNT3A strongly suppressed the midbrain marker LMX1A while WNT8A only partially suppressed it and WNT6 did not reduce its expression. WNT3 and WNT3A upregulated posterior markers including GBX2, CDX2, and HOX genes though thoracic markers such as HOXA7, HOXB8, and HOXC8. WNT8A induced GBX2 to a lesser degree and only induced more anterior HOX genes such as HOXA1, HOXA2, HOXB1, and HOXB2 (Figure 2D). Taken together these results suggest that WNT6 is only capable of suppressing the most anterior forebrain fates, WNT8A is capable of posteriorizing cells to mid or hindbrain fates but not further, while WNT3 and 3A exhibit strong posteriorizing activity.

### WNT ligands regulate overlapping transcriptional targets with different strength

To compare the global transcriptional effects of WNT ligands, we further examined the bulk RNA sequencing data from cells overexpressing WNT3, WNT3A, WNT8A, or WNT6 under neural ectoderm differentiation conditions (Figure 3A). Principal component analysis (PCA) revealed that samples clustered primarily by time point, with day 2 and day 6 separating along the first principal component (Figure 3B). At both time points, WNT3 and WNT3A clustered closely together, while WNT8A diverged moderately and WNT6 was the most distinct. WNT8A samples are consistently clustered closer to WNT3/3A than to WNT6.

**Figure 3:**
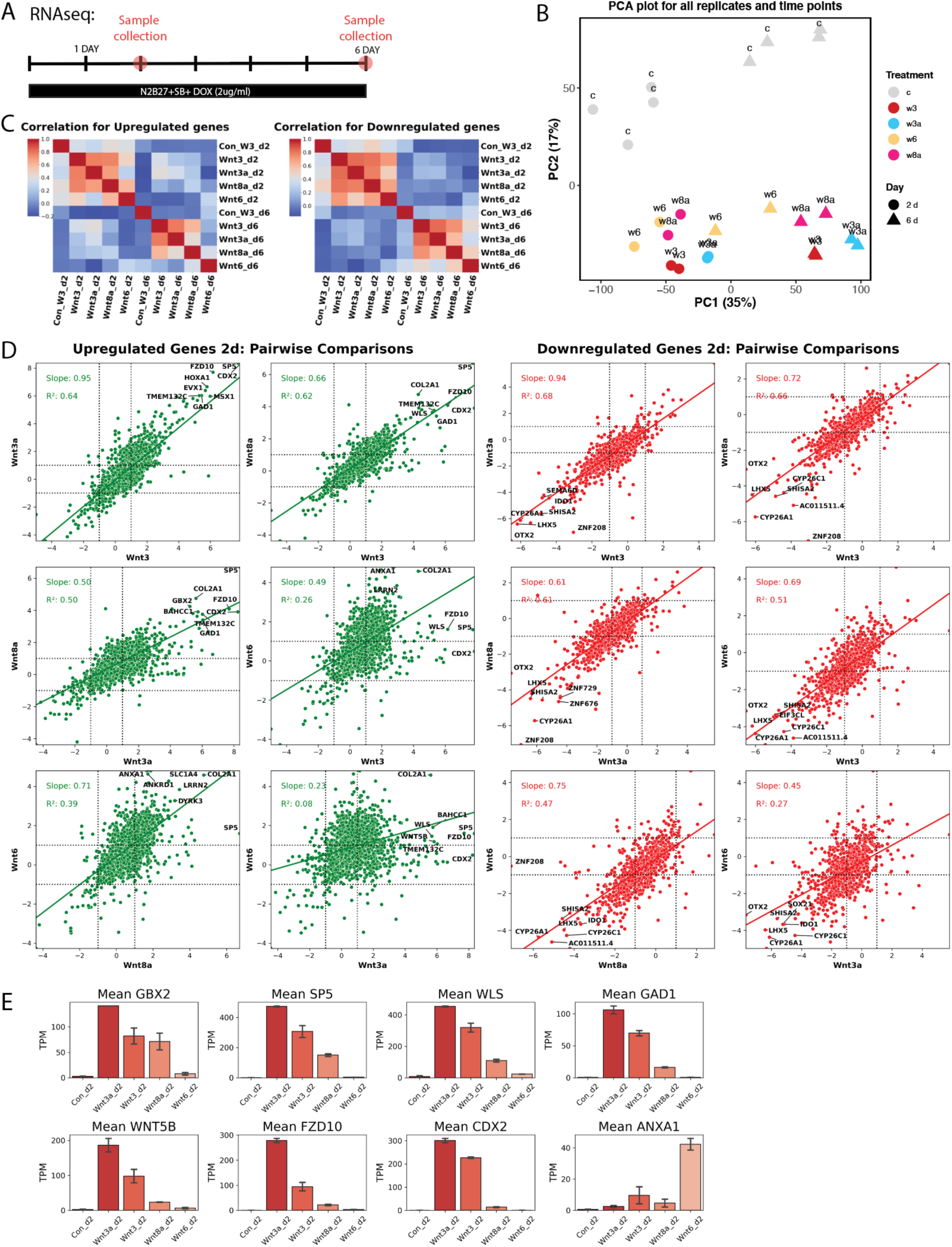
WNT ligands regulate partially overlapping sets of target genes with varying strength. (A). Schematic of differentiation protocol for the RNA-sequencing sample. (B). Principal component plot showing samples with different induced Wnt ligands for different treatment times. (C). Heatmaps of Pearson correlation for upregulated (left) and downregulated (right) genes (D). Scatter plots comparing expression induced by different Wnts. Best fit lines are shown and R^2^ and slope of line are indicated (E). Expression from RNA-sequencing data of selected gene targets differentially regulated between WNT ligands on day 2.

To compare expression profiles across ligands more directly, we focused on genes with substantial changes in expression(abs(log(2) FC) > 1 relative to control) and calculated pairwise Pearson correlations among WNT conditions (Figure 3C). WNT3, WNT3A, and WNT8A showed strong correlations in upregulated gene expression, while WNT6 was less correlated and aligned most closely with WNT8A. Correlations among downregulated genes were stronger and more consistent across all ligands, suggesting suppression of common targets. The trends between day 2 and day 6 were similar (Figure S5 A,B).

To further assess ligand-specific regulatory strength, we plotted pairwise gene expression comparisons for each ligand pair and fit linear models to both upregulated and downregulated genes. WNT3 and WNT3A regulated nearly identical gene sets with high *R*^2^ values (range: 0.60–0.66) and slopes near unity (0.90–0.92), indicating equivalent strength. WNT8A showed strong correlation with WNT3/3A but the slope of the best fit was ∼ 0.5, consistent with weaker induction of shared targets. WNT6 exhibited minimal correlation with WNT3/3A and only moderate correlation with WNT8A, suggesting it regulates a distinct transcriptional program (3D).

Despite these differences in upregulation, a substantial number of genes were downregulated by all four ligands in similar strengths. These included anterior neural markers (e.g., OTX2, LHX5, CYP26C1), WNT antagonists (SHISA2), and proliferation-associated genes (SEMA6D) 3D.

We used limma-voom for differential expression analysis, comparing each WNT condition to the control. Volcano plots revealed that on day 2, WNT3 and WNT3A had nearly identical effects, while WNT8A uniquely upregulated genes associated with fetal brain and optic neuron development (eg. ZNF512B, IRX6). WNT6 upregulated genes related to ion transport, enzymatic activity, and the cell cycle (eg. TMEM30B, PRSS1, TL3) (Figure S6 A).

By day 6, WNT3 and WNT3A strongly induced HOX gene expression, while WNT6 upregulated genes associated with cell adhesion and tight junctions (Figure 3F). Together, these analyses show that all four ligands downregulate a similar set of targets, while the targets upregulated by WNT6 differ from the other three ligands, and that the strength of regulation of shared targets varies between ligands.

### WNT8A strongly induces targets requiring moderate levels of Wnt signaling

Although WNT3 and WNT3A generally exhibited stronger signaling than WNT8A, RNA-seq analysis at day 6 identified several genes—SOX1, EGR2, and KLHL4—that were more strongly induced by WNT8A than by WNT3 or WNT3A (Figure 4A). Given that WNT8A otherwise activates many of the same targets as WNT3/3A but at lower levels, we hypothesized that these genes are preferentially induced at moderate WNT signaling strength and become suppressed at higher levels.

**Figure 4:**
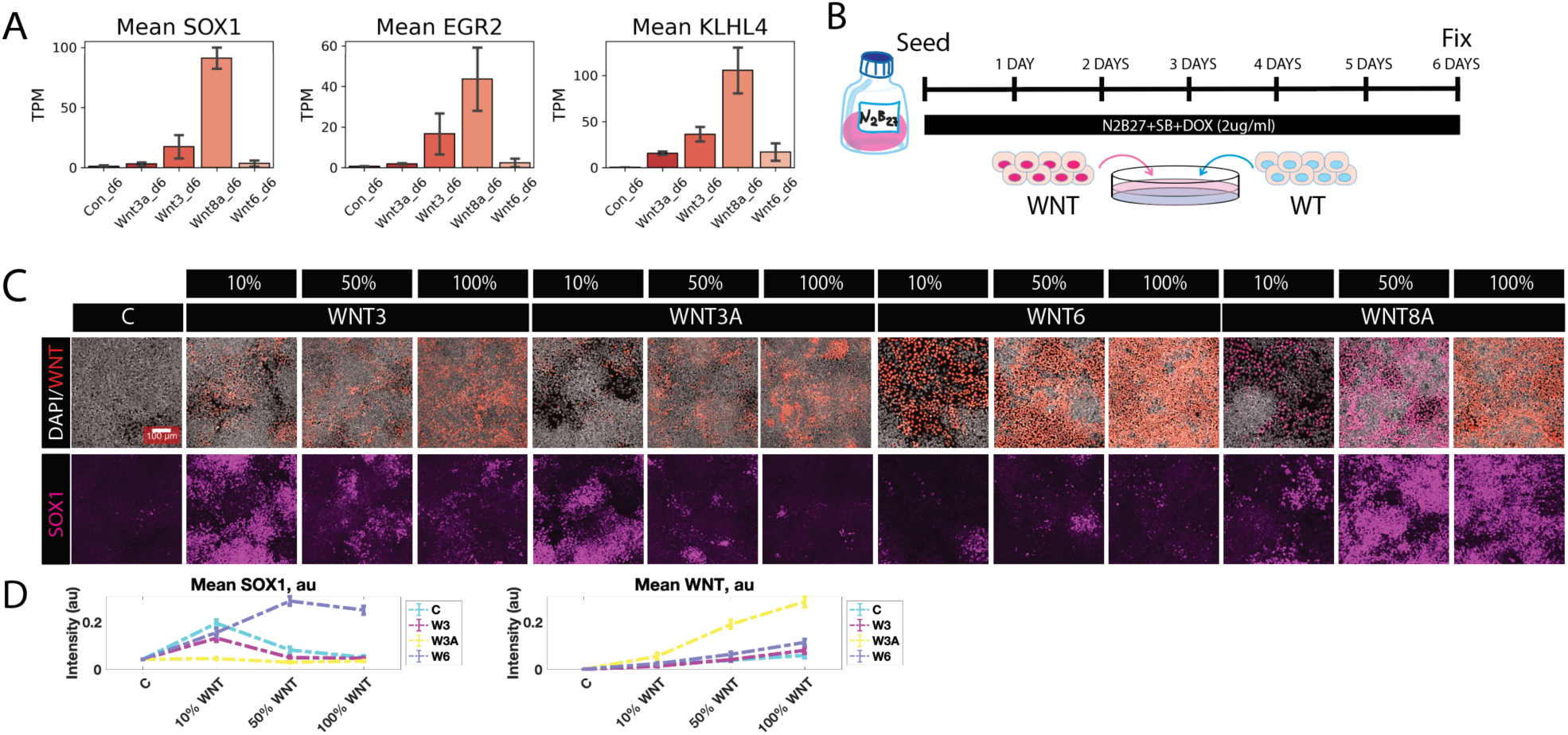
WNT8A strongly induces targets requiring intermediate WNT signaling levels. (A). RNA-seq identifies targets more strongly induced by WNT8A on day 6, including SOX1. (B). Schematic of the experiment. WNT-expressing cells were mixed with wild-type hPSCs at varying ratios. (C). Immunostaining of the indicated mixtures of cell lines for SOX1 (D). Dose-response curve of SOX1 expression as a function of the fraction of WNT-expressing cells.

To test this, we modulated WNT signaling using two complementary strategies. First, we mixed WNT-overexpressing cells with wild-type hESCs at defined ratios (10%, 50%, and 100% WNT-expressing cells), thereby modulated the total amount of WNT ligand (Figure 4B, Figure S7). SOX1 expression was absent in the 100% WNT3, WNT3A, and WNT6 conditions, but strongly induced in 100% WNT8A. As WNT3 and WNT3A levels were reduced by dilution, SOX1 expression increased, peaking at intermediate WNT levels. In contrast, SOX1 expression in WNT8A conditions declined as WNT8A was diluted, suggesting it no longer reached the activation threshold (Figure 4 C,D,E).

These results support a model in which SOX1 is induced only within a moderate window of WNT signaling: WNT8A naturally produces signaling levels that fall within this window, while WNT3 and WNT3A exceed it unless diluted. At high WNT activity, SOX1 is suppressed; at intermediate activity, it is activated (Figure 4 C). Notably, WNT6 failed to induce SOX1 at any level.

To independently validate these findings, we performed a titration of doxycycline in the WNT-overexpressing lines, rather than mixing cells. This yielded consistent results: SOX1 expression was maximally induced at intermediate doxycycline concentrations for WNT3 and WNT3A, while WNT8A required higher doxycycline to achieve comparable induction (Figure S7 C,D).

### WNT3/3A and WNT8A activate signaling several cell diameters beyond their expression domains

The diffusion of WNT ligands has been studied in a variety of contexts but direct comparisons of the range of action of different ligands are not available. We used a juxtaposition assay (described previously (Liu et al. 2022; Li et al. 2018)), to test the dispersion range of WNT ligands. We juxtaposed WNT expressing cells (“senders”) with wild-type cells (“receivers”) and treated the combined culture with doxycycline (Figure 5 A). After 6 days, cells were fixed and stained for markers of differentiation and WNT signaling activation. The experiment was performed under both pluripotent and neural induction conditions as described above. Under pluripotent conditions, we observed LEF1 expression in the WNT3/WNT3A sender cells and in the first two tiers of receiver cells adjacent to the sender cells. Notably, LEF1 expression was absent outside the sender domain in WNT8A-expressing cells. These results suggest that, under pluripotency conditions, WNT ligands exhibit minimal dispersal, extending between 0 and 2 cell diameters beyond the producing cells (Figure S8 A,B). In contrast, under neural ectoderm induction, LEF1 was more strongly induced and formed a gradient of expression which extended more than 10 cell diameters away from the sender cells (Figure 5 D,E). WNT8A induced lower LEF1 expression but with a similar spatial range. We also examined the expression of another direct WNT target, the secreted feedback inhibitor DKK1 (Figure 5 D, E). By day 5, senders expressed DKK1, and a stripe formed in the receivers with its expression overlapping with and extending beyond LEF1. How these expression patterns of DKK1 protein form, whether they involve diffusion, and whether DKK1 is important for restricting the activity of WNT targets such as LEF1 are interesting questions for future study. Another WNT inhibitor, AXIN2, formed a similar stripe-like pattern at the edges of the sender domain (Figure S8 C, D). These findings suggest the possibility of longer-range movement for WNT ligands during neural ectoderm patterning, highlighting the high degree of context dependency in WNT dispersion mechanisms.

**Figure 5:**
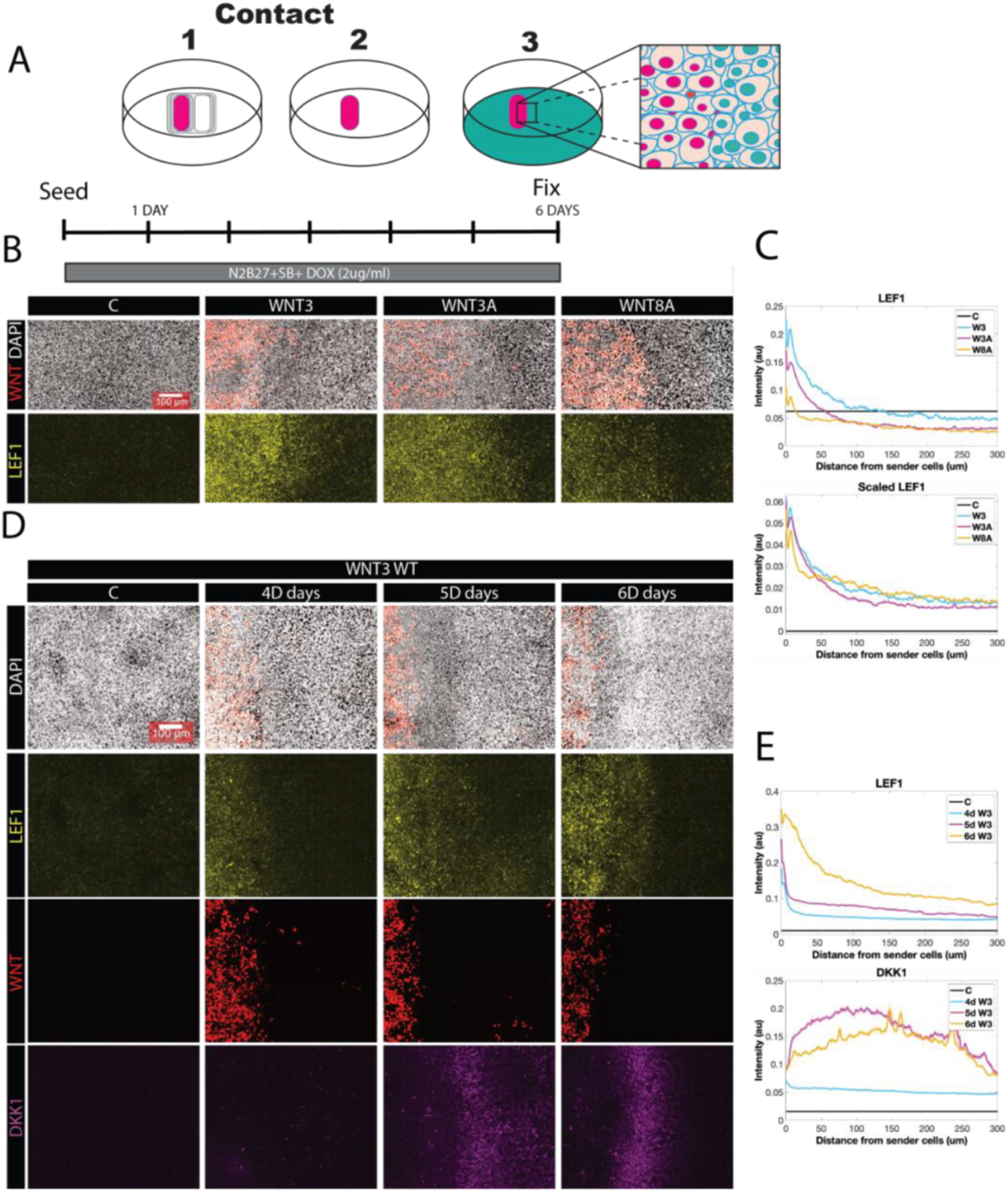
Spatial spread of Wnt ligands during ectodermal differentiation. (A). Schematic of the juxtaposition assay (B). Immunostaining of the indicated juxtaposition cultures for LEF1. (C) Quantification of the results in (B). The bottom panel shows the data with each curve rescaled so that the maximum of each curve is the same (D). Immunostaining for LEF1 and DKK1 in WNT3 juxtaposition cultures after 4, 5, or 6 days of juxtaposition. (E) Quantification of the results in (D).

### Localized WNT expression induces striped patterns of cell fates

We next examined the spatial patterns of cell fate that result from the juxtaposition experiments described above by analyzing markers of the anterior-posterior axis as well as of neural crest fates (Figure 6 A). We found that expression of either WNT3 or WNT3A suppressed OTX2 while inducing expression of SOX1 and HOXB1, consistent with a more posterior identity (Figure 6 B,C,D,E, Figure S9 A,B). The suppression of OTX2 extended approximately 6 cell diameters into the receiver cells and formed a sharp boundary between OTX2-positive and -negative domains even when the boundary between sender and receiver cells was not sharp (Figure 6B). WNT8A induced SOX1 but not HOXB1 and suppressed OTX2 more weakly. Its effects were restricted to areas immediately adjacent to the sender domain. WNT6 had only weak, cell-autonomous effects within the sender population itself (Figure 6 B, Figure S9 A,B).

**Figure 6:**
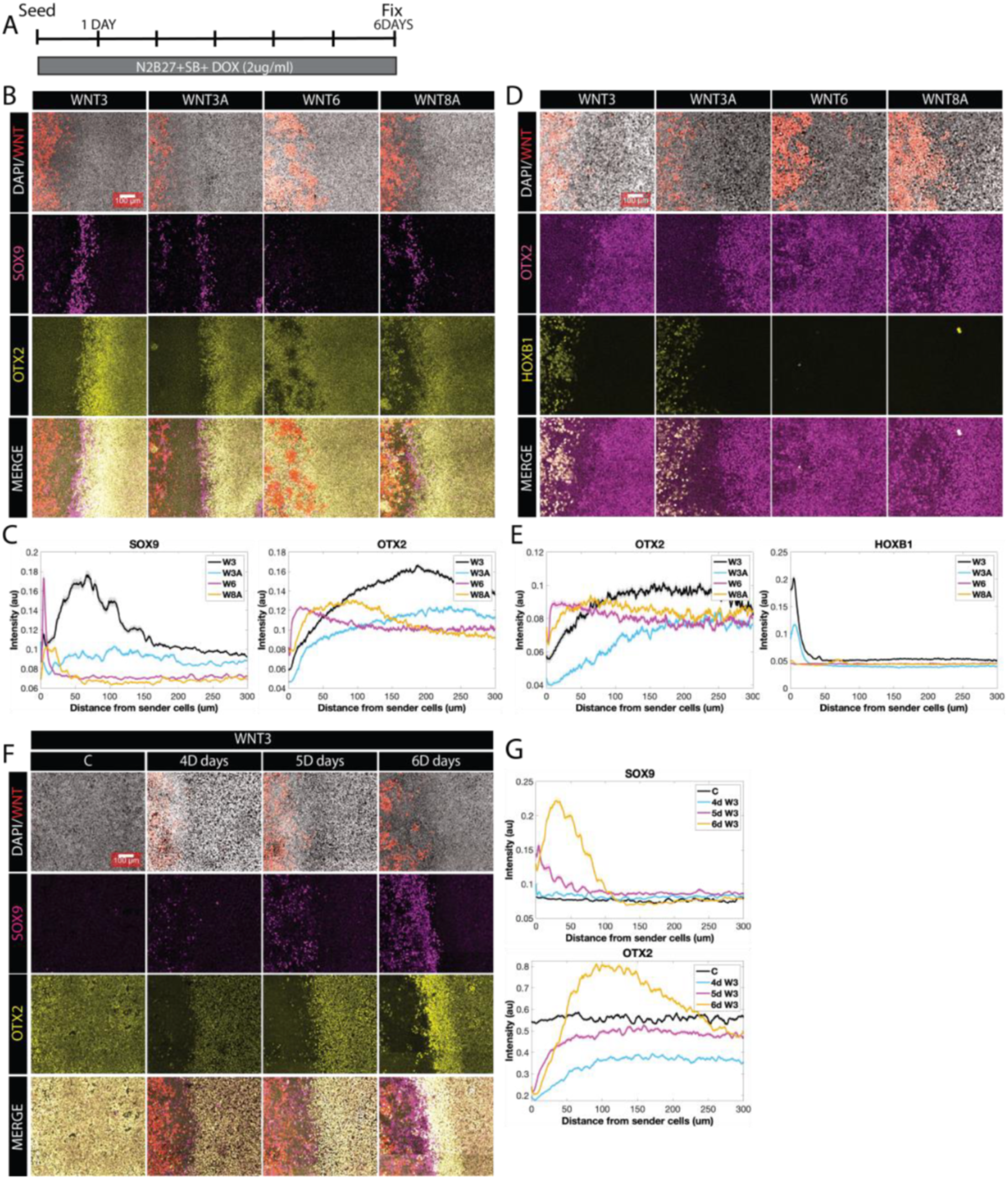
Localized sources of WNT ligands generate spatial patterns of ectodermal cell fates. (A). Schematic of 6-day juxtaposition assay under neural conditions. (B, D, F). Immunostaining of the juxtaposition cultures for markers of cell fates and AP patterning as indicated. Scale bars 100 um. (C,E,G) Quantification of the results in B,D, and F respectively.

In addition to their role in posteriorization, WNT ligands are implicated in neural crest cell (NCC) induction. NCC markers such as SOX9, PAX3, and AP2A are expressed downstream of WNT signaling (Chang and Hemmati-Brivanlou 1998; García-Castro et al. 2002; Gomez et al. 2019; Britton et al. 2019; Sutton et al. 2021)). In the juxtaposition assay, all four ligands induced some expression of these markers, but spatial patterns varied. WNT3 and WNT3A induced stripe-like expression domains of SOX9, PAX3, and AP2A peaking ≈ 2–3 cell diameters from the boundary (Figure 6 B, Figure S9 A,B). These domains were mutually exclusive with OTX2 and aligned with the steepest part of the WNT signaling gradient (Figure 6 B, F). WNT8A induced NCC markers with lower intensity and in a more compressed domain, while WNT6 induced only scattered expression restricted to the sender cells. We did not detect expression of the placodal marker SIX1 or the surface ectoderm marker ISL1, both of which require BMP activation for expression (Figure S10 B,C,F,G). Examining how these patterns emerged in time, we found that by day 4, OTX2 was already suppressed within the sender cells, while SOX9 expression was detected in scattered patches within the senders (Figure 6 F, G, Figure S9 D). On days 5 and 6, SOX9 expression coalesced into a stripe within the receivers and OTX2 expression strengthened at a greater distance from the senders, creating mutually exclusive domains of expression (Figure 6 B,F; Figure S9). These events occurred simultaneously with the strengthening and broadening of the LEF1 expression gradient and the emergence of DKK1 expression within the receivers. The relationship between these events is an interesting topic for future study (Figure S9 C,D,E,F, Figure S10 A).

In addition to WNT, both BMP and FGF/MEK signaling pathways have been implicated in neural crest induction. To assess their contributions in the context of spatial patterning, we repeated the WNT3 juxtaposition assay under BMP or MEK inhibition using Noggin or PD0325901, respectively (Figure 7A,B). BMP inhibition completely abolished SOX9 expression across the receiver domain while MEK inhibition led to a large reduction but not complete loss of SOX9 expression. We also examined OTX2 expression under these conditions. In all conditions examined, OTX2 is suppressed in receiver cells closest to the senders and OTX2 is expressed at a greater distance with a clear spatial boundary between the domains. In control WNT3 juxtaposition assays, OTX2 forms a narrow expression peak immediately adjacent to the cells with suppressed OTX2. At greater distances, the expression returns to the levels similar to the untreated control. However, under both BMP and MEK inhibition, after the gap, OTX2 expression remains uniformly high across the receiver domain (*Figure* 7B). This suggests that in the absence of these additional signals, WNT alone is insufficient to downregulate anterior identity, either due to reduced signaling activity or failure of the tissue to respond appropriately. This suggests that in the absence of these additional signals, WNT alone is insufficient to downregulate anterior identity, either due to reduced signaling activity or failure of the tissue to respond appropriately. These results highlight the need for combinatorially signaling through WNT, BMP, and MEK pathways to establish spatial patterns of cell fates.

**Figure 7:**
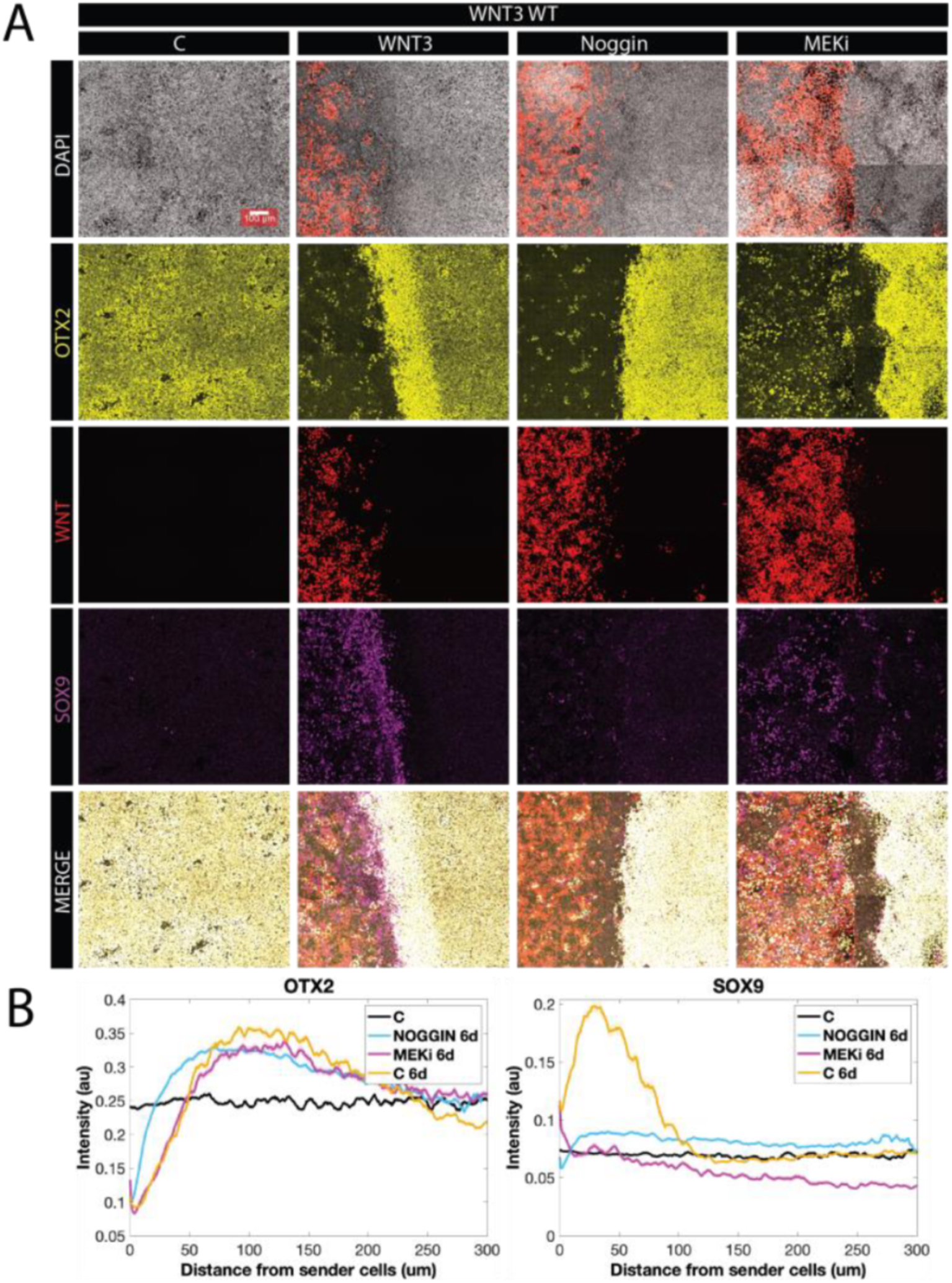
Dependence of Wnt-induced patterns on BMP and MEK signaling. (A). Juxtaposition assay with WNT3 expressing cells treated with Dox only or with Noggin or MEK-inhibitor added. (B). Quantification of the results in (A).

## Discussion

WNT ligands play crucial roles as regulators of cell fate and differentiation throughout development. While loss of function experiments in animal models have established distinct roles for a number of individual WNT ligands in early development, whether these result primarily from differences in expression patterns or intrinsic differences in ligand action has not been clarified. Here, we systematically compare the activity of WNT3, WNT3A, WNT6, and WNT8A in the context of early human cell fate decisions and patterning using hPSC-based models. We reveal that canonical WNT ligands, despite their structural similarity and common action through *β*-catenin signaling, exhibit distinct functional capacities. In particular, while WNT3 and WNT3A have very similar targets and strength, WNT8A influences a largely identical set of targets more weakly, while WNT6 represses a similar set of genes as the other WNT ligands but activates a distinct set. Our results highlight that different WNT ligands act across different spatial ranges to alter the transcription of overlapping, but distinct, sets of genes and influence cell fate decisions and patterning.

We examined the effects of WNT ligand stimulation in two different contexts, mesoderm induction from pluripotency and posteriorization during neural differentiation. WNT3/3A activated canonical targets more strongly during neural differentiation than in pluripotency, while a response to WNT8A was only seen in the neural context. WNT6 did not activate the same set of canonical target genes as the other ligands, but repressed similar genes, such as anterior markers, during neural differentiation. Thus, for the same ligand, signaling strength is context-dependent, potentially shaped by receptor expression, intracellular effectors, and inhibitory feedback mechanisms. In some cases, such as for WNT8A, the response is detectable in some contexts but not others.

We also assessed the spatial activity range of each WNT ligand. During mesoderm induction, WNT3/3A effects remained local, within 1-2 cell diameters from expressing cells. In contrast, WNT3, WNT3A, and WNT8A all induce LEF1 expression up to 8–10 cell diameters from the ligand-expressing source during neural differentiation, whereas the effects of WNT6 expression remain highly localized. These findings show that the range over which WNT ligands exert an effect is both ligand and context dependent. However, we cannot distinguish between direct extracellular movement of ligand and alternative mechanisms such as cell relay or contact-dependent signaling. Prior studies indicate that WNT3 may function as a short-range morphogen (Farin et al. 2016), while others suggest relay-like mechanisms (Kornberg 2017). Further experiments using tagged ligands or live imaging would be required to conclusively determine whether the observed range reflects ligand mobility or more indirect mechanisms of signal propagation. The use of WNT ligand knockout receiver cells could also help determine whether a relay mechanism involving induction of endogenous WNTs is involved.

Establishing the anteroposterior (AP) axis is a fundamental feature of early vertebrate development. WNTs are well-documented posteriorizing agents across species (McGrew et al. 1995; Kiecker and Niehrs 2001). Knockout studies in zebrafish, frogs, and mice have shown that loss of WNT3, WNT3A, or WNT5A leads to posterior truncation either as a result of loss of posterior identities, or as a failure of convergent extension movements (Amerongen and Berns 2006; Yamaguchi et al. 1999). Our findings show that WNT3 and WNT3A exhibit the strongest ability to activate posterior HOX genes while suppressing anterior markers such as OTX2 and PAX6. WNT8A shows weaker induction of only the most anterior HOX genes with no induction of more posterior genes, and WNT6 does not induce posterior markers, but does suppress anterior fates, consistent with our findings above that WNT6 has repressing but not activating targets in common with WNT3, 3A, and 8. The induction of more posterior fates by WNT3 and WNT3A also involves transient BRA expression on day 2. This likely reflects the need for cells of more posterior neural fates to first pass through a neuro-mesodermal progenitor (NMP) state before adopting spinal cord fates. These data reinforce the view that individual ligands possess distinct posteriorizing potential.

We found that when WNT ligands are expressed from a sender population of cells during neural differentiation, patterns emerge which involve stripes of both anterior-posterior markers such as OTX2 and HOX genes, as well as neural crest markers, such as PAX3 and SOX9. These neural crest markers would typically be found at a particular medial-lateral position along a broad range of the AP axis. This suggests that in this juxtaposition assay an axis is formed in the culture which conflates aspects of the AP and ML axes. It is an open question how WNT ligands can play roles in forming both of these orthogonal axes in vivo while in vitro a single composite axis forms. It would be interesting to understand the minimal requirements for developing two orthogonal WNT-dependent axes in a culture system.

Together, our findings challenge the assumption that canonical WNT ligands are functionally redundant. Despite established roles for all four ligands in the canonical WNT pathway, they differ in potency, diffusion range, and downstream transcriptional outcomes. These distinctions likely arise from receptor availability, extracellular modulation, and intrinsic biochemical differences. Our data argue for a ligand-centric view of WNT signaling, where individual WNTs should be studied as discrete actors rather than interchangeable parts of the pathway.

## Methods and Materials

All experiments in this study utilized the ESI-017 human embryonic stem cell (hESC) line, obtained from ESI BIO (RRID: CVCL-B854, XX), which is registered with the NIH. Karyotype and genomic integrity of the ESI-017 cells were verified by the manufacturer, and pluripotency markers (OCT4+, SOX2+, NANOG+) were consistently monitored throughout the study. Additionally, all cell lines were routinely screened for mycoplasma contamination and were confirmed to be negative. The research was conducted in accordance with the ISSCR guidelines for stem cell research and clinical translation that can be found at https://www.isscr.org/policy/guidelines-for-stem-cell-research-and-clinical-translation.

### Routine cell culture

All cells were cultured in chemically defined mTeSR1 medium (STEMCELL Technologies) in Matrigel-coated (Corning; 1:200 in DMEMF12) or GELTREX tissue culture dishes, maintained at 37°C with 5% CO2. Cell passage number did not exceed 60. Dispase (Fisher Scientific) was used for routine passaging, while Accutase (Corning) was used to prepare single-cell suspensions to seed for experiments. ROCK-inhibitor Y27672 (10 *μ*M; STEMCELL Technologies) was applied to support single-cell viability. For transfections with piggyBac plasmids containing WNT gene coding sequences, antibiotic selection with 4 *μ*g/ml Puromycin was employed for at least 2 days.

### Plasmids

Plasmids were constructed for this study as follows. The human WNT3, WNT3A, and WNT8A genes were amplified directly from hESC cDNA using standard PCR (OneTaq Quick-Load 2X MM, New England Biolabs, Inc.). Unique primers (Table S1) were designed to amplify and clone each gene into a plasmid backbone (Table S1). The WNT6 gene sequence was ordered from Genscript and similarly flanked by appropriate restriction enzyme sites. All WNT gene sequences included their stop codons and were validated using BLASTn. The TagRFP::NLS::T2A sequence was derived from the ePB-Bsd-CAG-RFP-NLS-T2A-Smad2 (AW-P6) plasmid and inserted into the ePiggyBac transposable element vector, epB-Bsd-TT-RFP (AW-P60). This vector, based on the pBSSK backbone, contains transposon-specific inverted terminal repeat sequences (ITRs) to ensure stable integration of the cassette between the ITRs. The sequence was inserted downstream of the doxycycline-induced TRE promoter, while Puromycin and the reverse tet transactivator (rTTA) were expressed under the human ubiquitin C (hUbC) promoter for stable selection (Table S1).

### DNA nucleofection and stable cell line establishment

DNA nucleofection was performed with the 4D-Nucleofector (Lonza) system according to the manufacturer’s protocol using the P3 Primary Cell 4D-Nucleofector Kit (Lonza). For ePiggyBac transposase mediated insertion, antibiotic selection started two days after nucleofection and lasted at least for 3 days. Antibiotics were not supplemented in medium for routine culture and were withdrawn at least 2–3 days prior to an experiment. A SH800S Cell Sorter (Sony) equipped with Cell Sorter software (version 2.2.4.5150) was used to select the 10% highest RFP expressing cells. After establishment, stable lines were checked for pluripotency markers, i.e., OCT4, SOX2 and NANOG expression, and found indistinguishable from WT ESI-017.

### Juxtaposition experiment

For juxtaposition experiments, hESCs overexpressing WNT were used as the sender cells, while wild-type hESCs (WT hESCs) acted as receiver cells. In the temporal analysis experiments, the sender cells were dissociated using Accutase, and 100,000 cells were re-suspended in 70 *μ*l of mTeSR1 media supplemented with the ROCK inhibitor Y27672 (10 *μ*M). These cells were then seeded into one well of a 2-well silicone insert (ibidi) with a 0.22 cm² growth area, which had been coated with Matrigel in a *μ*-Slide 8 Well (ibidi). The cells were incubated at 37°C for 1–2 hours to allow attachment to the culture surface, after which any unattached cells were washed away with PBS (without Ca/Mg) twice. The next day, the WT hESCs were also dissociated with Accutase. 500,000 cells were re-suspended in 250 *μ* l of mTeSR1 media supplemented with Y27672 (10 *μ*M), and seeded into the same well, juxtaposed with the sender cells. After 30-40 minutes, unattached receiver cells were washed out using PBS. Then, 250 *μ*l of either mTeSR1 or N2B27+SB431542 with 2 *μ*g/ml doxycycline (DOX) was added to the wells. The treatments were continued for the specified durations in each experiment, with daily media changes and re-addition of induction factors throughout, unless otherwise stated. Cells were fixed at designated time points for further analysis of various cell fate markers and signaling pathways.

### Immunofluorescence staining

Cells were washed twice with PBS and then fixed at room temperature using 4% paraformaldehyde for 25 min. After fixation, the samples were washed twice with PBS, followed by permeabilization and blocking using 3% donkey serum in PBST (1x PBS with 0.1% Triton X-100) for 30–60 minutes at room temperature. Primary antibodies were diluted in the blocking buffer as specified (Table S1). Cells were then incubated with the diluted primary antibody solution for 2 hours at room temperature or overnight at 4°C. After incubation, the cells were washed three times with PBST, each wash lasting 30 minutes. They were then treated with a secondary antibody solution diluted 1:500 (Table S1), which was supplemented with DAPI, and incubated for 1 hour at room temperature. The cells were then washed three times with PBST and placed in PBS (without Ca/Mg) prior to imaging.

### Fluorescent in situ hybridization

For dual smFISH/immunofluorescent labeling, the smFISH procedure (RNAscope™, Advanced Cell Diagnostics, Inc.) was performed first according to the RNAscope® Fluorescent Multiplex Assay. Cells were washed twice with PBS and then fixed at room temperature using 4% paraformaldehyde for 25 min. After fixation, the samples were washed twice with PBS, followed by gradual dehydration and rehydration according to the [RNAscope® Fluorescent Multiplex Assay manual chapter 3](http://fab.cba.mit.edu/classes/S63.21/class_site/pages//class_5/Supplemental%20Methods%20(TA)/320293_USM%20RNAscope%20Fluorescent%20Multiplex.pdf). The appropriate probe (RNAscope™ Probe- Hs-AXIN2-C2, Cat No. 400241-C2) was used to incubate samples for 2h at 40°C. Three amplification steps were followed until the final use of a fluorescent probe for the Far-Red channel. Finally, cells were washed with PBS (without Ca/Mg) and incubated with primary and secondary antibodies as specified in the immunofluorescence procedure above.

### Imaging

#### Fixed cell imaging

For all the fixed cell experiments, the images were taken with the 20x, 0.75 NA Objective on an Olympus FV3000 Laser Scanning microscope with a piezo ultrasound stage and the Olympus Software (with FV10-ASW 4.2 software). For standard cell culture experiments 4-6 images were taken per well. For juxtaposition experiments a 3x3 square grid was taken with the border area lying in the middle of the square.

#### Image analysis quantification/ code

Imaging experiments were conducted at least twice to ensure consistent results. Nuclear masks were generated based on the DAPI image using the trainable machine learning software ilastik (Berg et al. 2019). Images and masks were imported into MATLAB, and custom code was utilized to reduce background, to calculate fluorescent intensities, and for further analysis and visualization. The generated plots represent the mean and standard deviation of measurements taken from at least three fields of view. For juxtaposition experiments, a similar procedure was followed except a mask for one cell population was segmented using a fluorescent marker expressed in those cells (typically the sender cells). Protein expression in receiving cells was quantified as a function of their distance from producing cells. The distance was determined using the bwdist function in MATLAB, applied to the producing cell masks. Error bars in the resulting plots represent the SEM across multiple images at each time point.

### RNA-seq

#### Sample preparation

The treatments were conducted as outlined in the main text, using the following concentrations: doxycycline at 2 *μ*g/ml and SB at 10 *μ*M in N2B27 media for 2 and 6 days. Total RNA was extracted using the Invitrogen RNAqueous Micro Kit, and processed RNA was stored at -80∘C. RNA integrity was assessed via NanoDrop and agarose gel electrophoresis. Sequencing was performed by Novogene Co. on the Illumina NovaSeq 6000 platform, utilizing paired-end 150 sequencing. Quality control assessments were conducted by the sequencing provider. The raw sequencing data were processed using FastQC for quality checks and adapter trimming. Reads were pseudo-aligned to a reference transcriptome using Salmon, which performed transcript quantification in a Conda-managed environment to ensure reproducibility. Transcript abundance estimates were imported into R using the tximport package, and gene-level expression matrices were generated. Further downstream analysis was conducted in Python, where data were normalized to transcripts per million (TPM), scaled, and log-transformed to stabilize variance before exploratory analysis. Expression matrices were Z-score normalized per gene to facilitate comparisons across conditions. We used limma (‘version 3.58.1’) in R ("R version 4.3.2 (2023-10-31)") to perform Differential Gene Expression analysis because its voom transformation and empirical Bayes moderation robustly model the mean-variance relationship, enabling reliable statistical inference even with only two technical replicates per condition (Robinson et al. 2009; Law et al. 2014; Ritchie et al. 2015). Differential expression analysis identified genes with significant expression changes across experimental conditions - abs(log(2)FC)>1 and adjusted p-value < 0.05. Gene Ontology analysis was performed using GO.db in Python and cut offs of abs(log(2)FC)> 1 and adjusted p-value < 0.05. Visualization techniques, including scatter plots, heatmaps, venn diagrams and PCA plots were implemented using Seaborn and Matplotlib to explore expression trends. In the PCA plot, the overexpressed WNT- ligands were removed for ease of visualization. Volcano plots were generated in R using the limma and ggplot2 (‘version 3.5.1’) packages.

## Acknowledgments

We thank Lizhong Liu for helpful discussions regarding cloning and Cecilia Guerra for technical assistance. This work was supported by NIH grants R35GM149328 and R01HD112488 and NSF grant MCB-2135296 to AW.

## References

Barrott, J. J., Cash, G. M., Smith, A. P., Barrow, J. R., and Murtaugh, L. C. (2011). Deletion of mouse Porcn blocks Wnt ligand secretion and reveals an ectodermal etiology of human focal dermal hypoplasia/Goltz syndrome. Proceedings of the National Academy of Sciences of the United States of America, 108(31):12752–12757.540

Berg, S., Kutra, D., Kroeger, T., Straehle, C.N., Kausler, B.X., Haubold, C., Schiegg, M., Ales, J., Beier, T., Rudy, M., et al. (2019). Ilastik: Interactive Machine Learning for (Bio)Image Analysis. Nat. Methods 16, 1226–1232.

Biechele, S., Cockburn, K., Lanner, F., Cox, B. J., and Rossant, J. (2013). Porcn-dependent Wnt signaling is not required prior to mouse gastrulation. Development, 140(14):2961–2971.542

Boutros, M. and Niehrs, C. (2016). Sticking Around: Short-Range Activity of Wnt Ligands. Developmental Cell, 36(5):485–486.

Britton, G., Heemskerk, I., Hodge, R., Qutub, A. A., and Warmflash, A. (2019). A novel self-organizing embryonic stem cell system reveals signaling logic underlying the patterning of human ectoderm. preprint 20, Developmental Biology.

Chang, C. and Hemmati-Brivanlou, A. (1998). Cell fate determination in embryonic ectoderm. Journal of Neurobiology, 36(2):128–151.549

Chhabra, S., Liu, L., Goh, R., Kong, X., Warmflash, A., Kong, X., and Warmflash, A. (2019). Dissecting the dynamics of signaling events in the BMP, WNT, and NODAL cascade during self-organized fate patterning in human gastruloids. PLOS Biology, 17(10):e3000498.552

Christian, J. L., McMahon, J. A., McMahon, A. P., and Moon, R. T. (1991). Xwnt-8, a Xenopus Wnt-1 /int-1-related gene reponsive to mesoderm-inducing growth factors, may play a role in ventral mesodermal patterning during embryogenesis. Development, 111(4):1045–1055.555

Clevers, H. (2006). Wnt/*β*-Catenin Signaling in Development and Disease. Cell, 127(3):469–480.

Cunningham, T. J., Kumar, S., Yamaguchi, T. P., and Duester, G. (2015). Wnt8a and Wnt3a cooperate in the axial stem cell niche to promote mammalian body axis extension. Developmental dynamics: an official publication of the American Association of Anatomists, 244(6):797–807.

Dijksterhuis, J. P., Baljinnyam, B., Stanger, K., Sercan, H. O., Ji, Y., Andres, O., Rubin, J. S., Hannoush, R. N., and Schulte, G. (2015). Systematic mapping of WNT-FZD protein interactions reveals functional selectivity by distinct WNT-FZD pairs. Journal of Biological Chemistry, 290(11):6789–6798.562

Farin, H. F., Jordens, I., Mosa, M. H., Basak, O., Korving, J., Tauriello, D. V. F., de Punder, K., Angers, S., Peters, P. J., Maurice, M. M., and Clevers, H. (2016). Visualization of a short-range Wnt gradient in the intestinal stem-cell niche. Nature, 530(7590):340–343.565

Ferretti, E. and Hadjantonakis, A. K. (2019). Mesoderm specification and diversification: from single cells to emergent tissues. Current Opinion in Cell Biology, 61:110–116.567

Garcıa-Castro, M. I., Marcelle, C., Bronner-Fraser, M., García-Castro, M. I., Marcelle, C., and Bronner-Fraser, M. (2002). Ectodermal Wnt Function as a Neural Crest Inducer. Science, 297(5582):848–851.569

Gomez, G. A., Prasad, M. S., Sandhu, N., Shelar, P. B., Leung, A. W., and García-Castro, M. I. (2019). Human neural crest induction by temporal modulation of WNT activation. Developmental Biology, 449(2):99–106.571

Green, D., Whitener, A. E., Mohanty, S., and Lekven, A. C. (2015). Vertebrate nervous system posteriorization: Grading the function of Wnt signaling. Developmental Dynamics, 244(3):507–512.

Haegel, H., Larue, L., Ohsugi, M., Fedorov, L., Herrenknecht, K., and Kemler, R. (1995). Lack of *β*-catenin affects mouse development at gastrulation. Development, 121(11):3529–3537.575

Kiecker, C. and Niehrs, C. (2001). A morphogen gradient of Wnt/*β*-catenin signalling regulates anteroposterior neural patterning in Xenopus. Development, 128(21):4189–4201.577

Kornberg, T. B. (2017). Distributing signaling proteins in space and time: the province of cytonemes. Current Opinion in Genetics Development, 45(10):22–27.579

Law, C.W., Chen, Y., Shi, W., and Smyth, G.K. (2014). Voom: Precision weights unlock linear model analysis tools for RNA-seq read counts. Genome Biol. 15, 1–17.

Li, P., Markson, J. S., Wang, S., Chen, S., Vachharajani, V., and Elowitz, M. B. (2018). Morphogen gradient reconstitution reveals Hedgehog pathway design principles. Science, 360(6388):543–548.581

Lippmann, E. S., E.williams, C., Ruhl, D. A., Estevez-Silva, M. C., Chapman, E. R., Coon, J. J., and Ashton, R. S. (2015). Deterministic HOX patterning in human pluripotent stem cell-derived neuroectoderm. Stem Cell Reports, 4(4):632–644.584

Liu, L., Nemashkalo, A., Rezende, L., Jung, J. Y., Chhabra, S., Guerra, M. C., Heemskerk, I., and Warmflash, A. (2022). Nodal is a short-range morphogen with activity that spreads through a relay mechanism in human gastruloids. Nature Communications, 13(1):1–12.587

Liu, P., Wakamiya, M., Shea, M. J., Albrecht, U., Behringer, R. R., and Bradley, A. (1999). Requirement for Wnt3 in vertebrate axis formation. Nature Genetics, 22(4):361–365.589

Logan, C. Y. and Nusse, R. (2004). The Wnt signaling pathway in development and disease. Annual Review of Cell and Developmental Biology, 20(1):781–810.591

McGrew, L. L., Lai, C. J., and Moon, R. T. (1995). Specification of the Anteroposterior Neural Axis through. Synergistic Interaction of the Wnt Signaling Cascade with noggin and follistatin. Developmental Biology, 172(1):337–342.

Mikels, A. J. and Nusse, R. (2006). Purified Wnt5a protein activates or inhibits *β*-catenin-TCF signaling depending on receptor context. PLoS Biology, 4(4):570–582.596

Minn, K. T., Dietmann, S., Waye, S. E., Morris, S. A., and Solnica-Krezel, L. (2021). Gene expression dynamics underlying cell fate emergence in 2D micropatterned human embryonic stem cell gastruloids. Stem Cell Reports, 16(5):1210–1227.599

Niehrs, C. (2012). The complex world of WNT receptor signalling. Nature Reviews Molecular Cell Biology, 13(12):767–779.601

Niehrs, C. (2022). The role of Xenopus developmental biology in unraveling Wnt signalling and antero-posterior axis formation. Developmental Biology, 482(September 2021):1–6.603

Nusse, R. and Clevers, H. (2017). Wnt/*β*-Catenin Signaling, Disease, and Emerging Therapeutic Modalities. Cell, 169(6):985–999.605

Ritchie, M.E., Phipson, B., Wu, D., Hu, Y., Law, C.W., Shi, W., and Smyth, G.K. (2015). Limma powers differential expression analyses for RNA-sequencing and microarray studies. Nucleic Acids Res. 43, e47.

Robinson, M.D., McCarthy, D.J., and Smyth, G.K. (2009). edgeR: A Bioconductor package for differential expression analysis of digital gene expression data. Bioinformatics 26, 139–140.

Sutton, G., Kelsh, R. N., and Scholpp, S. (2021). Review: The Role of Wnt/*β*-Catenin Signalling in Neural Crest Development in Zebrafish. Frontiers in Cell and Developmental Biology, 9(November):1–15.607

Takada, S., Stark, K. L., Shea, M. J., Vassileva, G., McMahon, J. A., and McMahon, A. P. (1994). Wnt-3a regulates somite and tailbud formation in the mouse embryo. Genes and Development, 8(2):174–189.609

Tanaka, S. S., Kojima, Y., Yamaguchi, Y. L., Nishinakamura, R., and Tam, P. P. L. (2011). Impact of WNT signaling on tissue lineage differentiation in the early mouse embryo. Development, Growth and Differentiation, 53(7):843–856.

Tao, Y., Mis, M., Blazer, L., Ustav Jnr, M., Steinhart, Z., Chidiac, R., Kubarakos, E., O’Brien, S., Wang, X., Jarvik, N., Patel, N., Adams, J., Moffat, J., Angers, S., and Sidhu, S. S. (2019). Tailored tetravalent antibodies potently and specifically activate wnt/frizzled pathways in cells, organoids and mice. eLife, 8:1–16.

Tyser, R. C., Mahammadov, E., Nakanoh, S., Vallier, L., Scialdone, A., and Srinivas, S. (2020). A spatially resolved single cell atlas of human gastrulation.

Vallier, L. (2005). Activin/Nodal and FGF pathways cooperate to maintain pluripotency of human embryonic stem cells. Journal of Cell Science, 118(19):4495–4509.

van Amerongen, R. and Berns, A. (2006). Knockout mouse models to study Wnt signal transduction. Trends in Genetics, 22(12):678–689.

van Amerongen, R. and Nusse, R. (2009). Towards an integrated view of Wnt signaling in development. Development, 136(19):3205–3214.

Voloshanenko, O., Gmach, P., Winter, J., Kranz, D., and Boutros, M. (2017). Mapping of Wnt-Frizzled interactions by multiplex CRISPR targeting of receptor gene families. FASEB Journal, 31(11):4832–4844.

Warmflash, A., Sorre, B., Etoc, F., Siggia, E. D., and Brivanlou, A. H. (2014). A method to recapitulate early embryonic spatial patterning in human embryonic stem cells. Nature Methods, 11(8):847–854.

Wei, M., Zhang, C., Tian, Y., Du, X., Wang, Q., and Zhao, H. (2020). Expression and Function of WNT6: From Development to Disease. Frontiers in Cell and Developmental Biology, 8(December):1–10.629

Wylie, A. D., Fleming, J. A. G., Whitener, A. E., and Lekven, A. C. (2014). Post-transcriptional regulation of wnt8a is essential to zebrafish axis development. Developmental Biology, 386(1):53–63.631

Yamaguchi, T., Bradley, A., McMahon, A., and Jones, S. (1999). A Wnt5a pathway underlies outgrowth of multiple structures in the vertebrate embryo. Development, 126(6):1211–1223.633

Yamaguchi, T. P. (2001). Heads or tails: Wnts and anterior–posterior patterning. Current Biology, 11(17):R713–R724.

