## Supplementary Table and Figures for "Comparison between the activities of canonical Wnt ligands in human pluripotent stem cell differentiation"

### Supplement Figures and Tables

**Table S1. Key resources.**

*Antibodies & FISH probes*

| Antibodies | Supplier | Catalog Number |
|---|---|---|
| G BRACHYURY (1:300) | R&D Systems | Cat#AF2085 |
| M CDX2 (1:50) | Biogenex | Cat#MU392A |
| G HAND1 (1:200) | R&D Systems | Cat#AF3168 |
| M ISL1 (1:50) | Developmental Studies Hybridoma Bank | Cat#39.4D5 |
| R SOX2 (1:300) | Cell Signaling Technologies | Cat#5024S |
| R OCT3/4 (1:300) | BD Biosciences | Cat#611203 |
| G NANOG (1:100) | R&D Systems | Cat#AF1997 |
| G SOX17 (1:200) | R&D Systems | Cat#AF1924 |
| R pSMAD1 (1:200) | Cell Signaling Technologies | Cat#13820 |
| R SMAD2/3 (1:100) | BD Biosciences | Cat#610842 |
| M TFAP2A (1:50) | Developmental Studies Hybridoma Bank | Cat#AB_528084 |
| R PAX6 (1:300) | Biolegend | Cat#901301 |
| M PAX3 (1:50) | Developmental Studies Hybridoma Bank | Cat#AB_2722285 |
| G SOX9 (1:300) | R&D Systems | Cat#AF3075 |
| R OTX2 (1:300) | Abcam | Cat#ab21990 |
| G SOX1 (1:300) | R&D Systems | Cat#AF3369 |
| G SIX1 (1:300) | Cell Signaling Technology | Cat#12891S |
| G DKK1 (1:300) | R&D Systems | Cat#AF1096-SP |
| R LEF1 (1:300) | Cell Signalling Technology | Cat#2230s |
| Probe- Hs-AXIN2-C2 | RNAscope™ , ACD Inc. | Cat#400241-C2 |

*Bacterial Strain*

| Bacterial Strain | Supplier | Catalog Number |
|---|---|---|
| NEB®10-beta Competent E.coli (High Efficiency) | New England Biolabs (NEB) | Cat#C3019H |
| NEB®5-alpha Competent E.coli (High Efficiency) | New England Biolabs | Cat#C2987H |
| NEB® Stable Competent E.coli (High Efficiency) | New England Biolabs (NEB) | Cat#C3040H |

*Chemicals, Peptides, and Recombinant Proteins*

| Reagent | Supplier | Catalog Number |
|---|---|---|
| SB431542 | Stemgent | Cat#04-0010-05 |
| BMP4 | Fisher Scientific | Cat#314BP050 |
| CHIR 99021 | Fisher Scientific | Cat#HY-10182 |
| ROCK inhibitor Y-27632 | Fisher Scientific | Cat#50-175-998 |
| doxycycline HYCLATE | Sigma-Aldrich | Cat#D9891-1G |
| cOmpleteTM Lysis-M | MedChem Express | Cat#04719956001 |
| cloneR | Sigma-Aldrich | Cat#05889 |
| DAPI (4,6-diamidino-2- phenylindole, dihydrochloride) | STEM CELL Technologies | Cat#D1306 |
| DL-Dithiothreitol | Sigma-Aldrich | Cat# D9163-5G |
| IWP2 | Stemgent | Cat#04-0034 |
| mTeSR1 | STEM CELL Technologies | Cat#85875 |
| Dulbecco's PBS Without calcium and magnesium | Caisson Labs | Cat# PBL01-6X500ML |
| Puromycin | Fisher Scientific | Cat#A1113803 |
| Geltrex Ldev free heSC qual. | Life Technologies | Cat#A1413302 |
| Matrigel hESC-qualified Matrix | Corning | Cat#354277 |
| B-27 Supplement 50x, minus VitA (For N2B27media) | Fisher Scientific | Cat#12587010 |
| N2-Supplement (for N2B27 media) | Fisher Scientific | Cat#17502048 |
| Glutamax I 100x (200mM) | Life Technologies | Cat#35050061 |
| Dulbecco's Modification of Eagle's Medium (DMEM) 4.5 g/L glucose, L-glutamine, and sodium pyruvate | VWR | Cat#45000-304 |
| Neurobasal Medium (L-glutamine) | Fisher Scientific | Cat#21103-049 |

*Table S2. Critical Commercial Assays*

| Assay | Supplier | Catalog Number |
|---|---|---|
| DNeasy Blood and Tissue Kit | Qiagen | Cat#69504 |
| Miniprep Plasmid Kits, IBI Scientific, Hi-Speed Mini Plasmid Kit, Kit Size=300 Preps | IBI Scientific | Cat#IB47102 |
| HiSpeed Plasmid Midi Kit | Qiagen | Cat#12643 |

| Assay | Supplier | Catalog Number |
|---|---|---|
| InvitrogenTM TOPOTM TA CloningTM Kit for Sequencing, without competent cells | Fisher Scientific | Cat#450030 |
| P3 Primary Cell 4D-Nucleofector® X Kit L | Lonza | Cat# V4XP-3024 |
| Gel extraction, DNA Recovery Kit, Zymoclean™(uncapped columns), 200 preps | Fisher Scientific | Cat#D4002 |
| RNAqueous®-Micro Total RNA Isolation Kit | Fisher Scientific | Cat#AM1931 |
| SuperScript Vilo cDNA Synthesis Kit | Fisher Scientific | Cat#11754-050 |
| RNA Clean & Concentrator™-5 | Zymo Research | Cat#R1015 |
| OneTaq Quick-Load 2X Master Mix with Standard Buffer – 100 reactions (50 µl vol) | New England Biolabs, Inc. | Cat#50-439-4 |

*Experimental Models: Cell Lines*

| Cell lines | Supplier | Catalog Number |
|---|---|---|
| ESI-017 | ESI BIO [16] | RRID:CVCL B854 |

*Primers for WNT cloning*

| Primer name | Supplier | Sequence |
|---|---|---|
| 6.1 fixGapNewfwd correct | Integrated DNA Technologies | 5' GGAGAATTCGAGCTCGGTACCCGGtttaattaatctcgacggtatcggttaacg 3' |
| 6.1 fixGapNewrev correct | Integrated DNA Technologies | 5' GGAGCAGCCCGAGCAGGTGGGGCTCCATGGATCCAGGGCCGGGATTC 3' |
| 2. CONTROL wnt3 Fwd BAMHI | Integrated DNA Technologies | 5' CTAGTGGATCCAAGAAATGGAGCCCCACCTGCTCGGGCTG 3' |
| 2. control WNT3 Rev NOTI | Integrated DNA Technologies | 5' GCGGCCGCATCTCCCTACTTGCAGGTGTGCACGTCGTAGATGCG 3' |
| Fwd cDNA WNT3A | Integrated DNA | 5' aatcccggccctGGATCCATGGCCCCACTCGGATACTTCTTAC 3' |

| Primer name | Supplier | Sequence |
|---|---|---|
| | Technologies | |
| Rev cDNA WNT3A | Integrated DNA Technologies | 5' gattatgatctagagtcgcCTACTTGCAGGTGTGCACGTCGTAG 3' |
| Fwd WNT6gen 1 | Integrated DNA Technologies | 5' gacgtggaggagaatcccggccctGGATCCATGCTGCCGCCCTTACCC 3' |
| Rev WNT6gen 1 | Integrated DNA Technologies | 5' gtatggctgattatgatctagagtcgcggccGCTCACAGGCAGAGGCTGAG 3' |
| WNT8A 6.1 FWD BamHI new | Integrated DNA Technologies | 5' cggtgacgtggaggagaatcccggccctGGATCCATGGGGAACCTGTTTATGCTCTG 3' |
| WNT8A-6.1 REV NotI Over | Integrated DNA Technologies | 5' CCcgcggccgcTCAGGCACTGCCCTTACC 3' |

*Primers for Sequencing*

| Primer name | Supplier | Sequence |
|---|---|---|
| Seq epB-Bsd-CAG-FWD 1 | Integrated DNA Technologies | 5' TACAGCTCCTGGGCAACGTGCT 3' |
| Seq ePB-Bsd-CAG-REV3 | Integrated DNA Technologies | 5' CAACAACAATTGCATTCATTTTATG 3' |
| Seq F RFP control | Integrated DNA Technologies | 5' caaggaggccgacaaagagacc 3' |
| Seq puroDownst F | Integrated DNA Technologies | 5' cgaggcgcaccgtgggcttg 3' |
| Seq test AWP60 Fwd | Integrated DNA Technologies | 5' GTTCTAAGGCCGAGTCTTATG 3' |

*Software and Algorithms*

| Software | Supplier |
|---|---|
| Benchling | https://benchling.com/ |

| Software | Supplier |
| --- | --- |
| ilastik | http://ilastik.org/ |
| MATLAB | https://www.mathworks.com/products/matlab.html |
| MATLAB scripts for quantifying, analyzing data and running simulations | https://github.com/eleanarizou/PaperUNO |
| R, R studio, R version 4.3.2 (2023-10-31) | https://www.R-project.org/ |
| R scripts for quantifying, analyzing RNA sequencing data | https://github.com/eleanarizou/PaperUNO |
| PYTHON, jupyter lab | https://www.python.org |
| python scripts for quantifying, analyzing RNA sequencing data | https://github.com/eleanarizou/PaperUNO |

*Recombinant DNA and Plasmids*

| Plasmid | Supplier | Sequence |
| --- | --- | --- |
| pSpCas9(BB)-2A-Puro (PX459) | addgene, Cat#48139 | - |
| GENEWIZ -WNT6 | Genewiz, | - |
| epB-Bsd-TT-RFP | (AWP60) | - |
| ePB-Bsd-CAG-RFP-NLS-T2A-Smad2 | (AWP6) | - |
| 6. 1. ePB-Puro-TT-RFP-NLS-T2A-WNT3 | EleanaR | ’WNT3’ |
| 6. 1. ePB-Puro-TT-RFP-NLS-T2A-WNT3A | EleanaR | ’WNT3A’ |
| 6.1 ePB-Puro-TT-RFP-NLS-T2A-WNT6 | EleanaR | ’WNT6’ |
| 6.1. ePB-Puro-TT-RFP-NLS-T2A-WNT8Acon | EleanaR | ’WNT8A’ |

*Cell culture reagents*

| Reagent | Supplier | Catalog Number |
| --- | --- | --- |
| Geltrex Ldev free heSC qual. | Life Technologies | Cat#A1413302 |
| Matrigel hESC-qualified Matrix | Corning | Cat#354277 |
| B-27 Supplement 50x, minus VitA (For N2B27media) | Fisher Scientific | Cat#12587010 |
| N2-Supplement (for N2B27 media) | Fisher Scientific | Cat#17502048 |
| Glutamax I 100x (200mM) | Life Technologies | Cat#35050061 |

| Reagent | Supplier | Catalog Number |
|---|---|---|
| Dulbecco's Modification of Eagle's Medium (DMEM) 4.5 g/L glucose, L-glutamine, and sodium pyruvate | VWR | Cat#45000-304 |
| Neurobasal Medium (L-glutamine) | Fisher Scientific | Cat#21103-049 |

*Dishes Plates*

| Dish | Supplier | Sequence |
|---|---|---|
| $\mu$-Slide 18 Well dishes | ibidi | Cat#81816 |
| $\mu$-Slide 8 Well High Glass Bottom | ibidi | Cat#80807 |
| Corning® 96-well Flat Clear Bottom Black Polystyrene TC-treated Microplates, Individually Wrapped, with Lid, Sterile | Fisher Scientific | Cat#07-200-565 |
| 35mm Culture Dish [15366] Thermo Scientific Nunc Dishes, Cell Culture/Petri - with Airvent, 500/Cs | Fisher Scientific | Cat#12-565-91 |
| 12well dish ThermoScient 150628 Nunclon Delta MultiDishes | Fisher Scientific | Cat#12-565-321 |

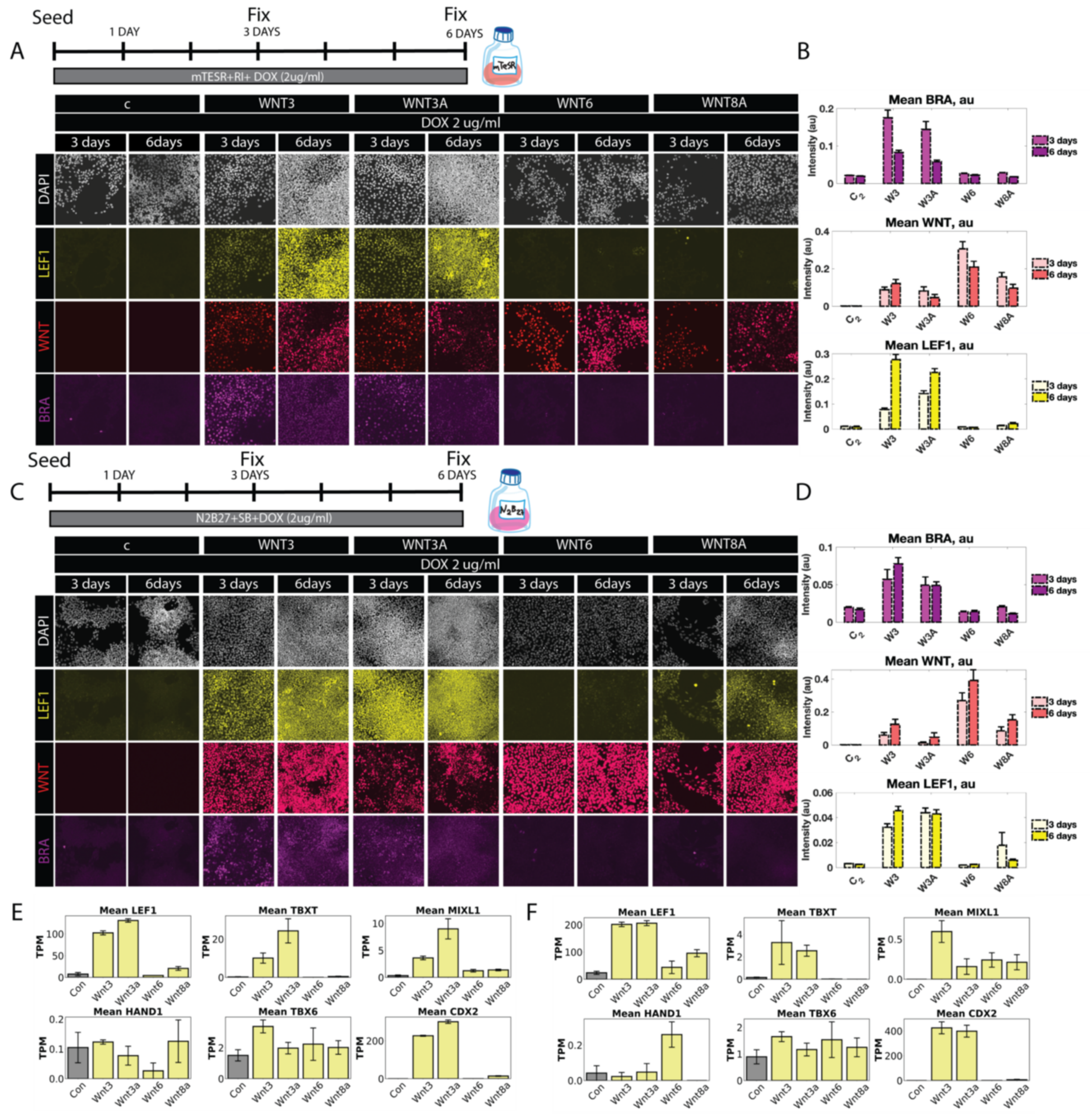


**Figure S1. WNT3 and WNT3A induce mesodermal differentiation but WNT6 and WNT8A do not.**

(A, C) Immunostaining of the indicated cell lines for LEF1 and BRA expression with dox added for the indicated time to either pluripotency media (A) or neuroectoderm differentiation media (C).

(B, D). Quantification of the results in A (B) and C (D).

(E, F). Bulk RNA-seq analysis of mesoderm marker expression at day 2 (E) and day 6 (F) during neuroectoderm differentiation.

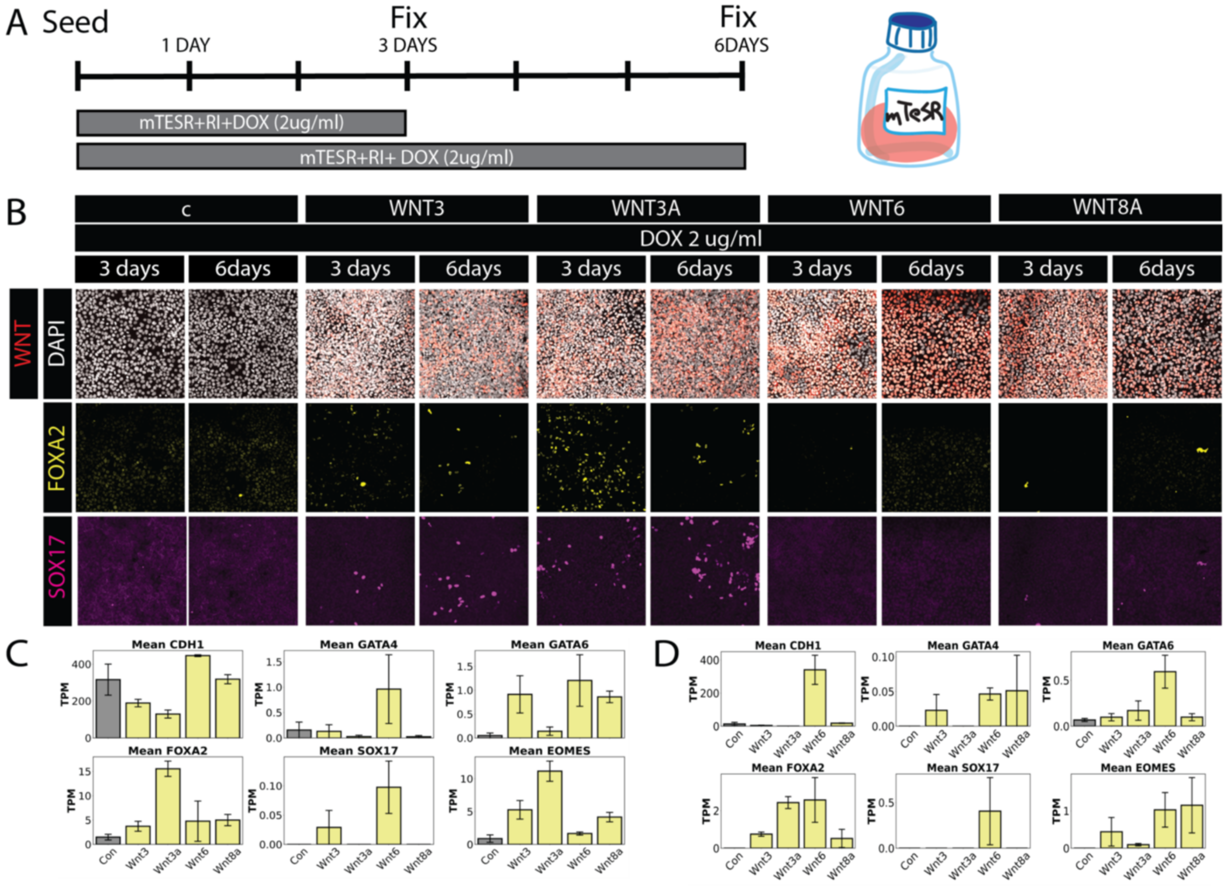


**Figure S2. WNT ligands do not significantly induce endodermal markers.**

(A). Schematic of the experiment.

(B). Immunostaining of the indicated cell lines for SOX17 and FOXA2 expression with dox added for the indicated time

(C,D). Bulk RNA-seq of endoderm markers at 2 (C) or 6 (D) days in neural ectoderm differentiation.

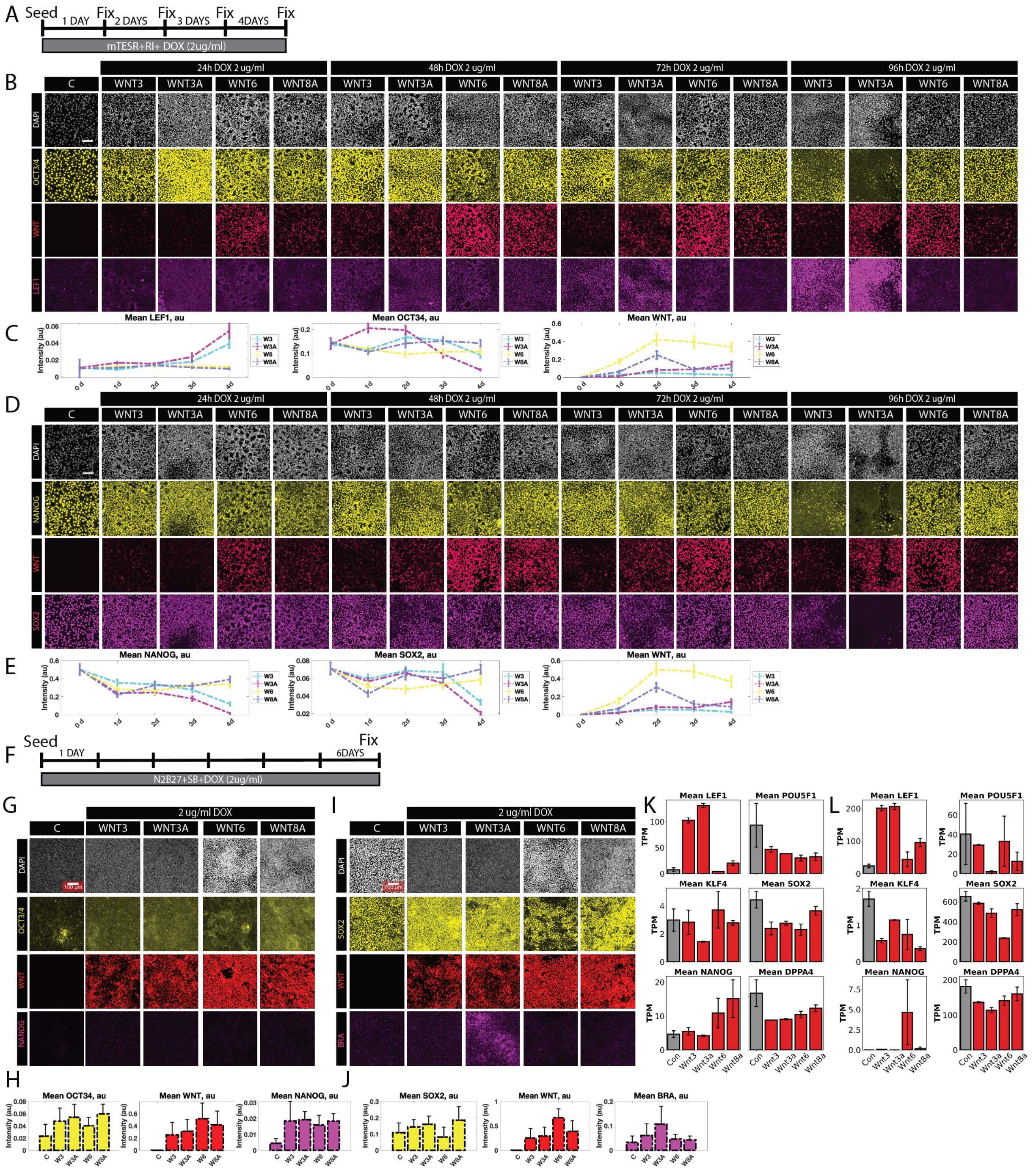

**Figure S3. WNT3/3A downregulate pluripotency markers over time, while WNT6 and WNT8A show weaker effects.**

(A) Schematic of the experimental conditions. (B, D) Immunostaining for OCT4 (B) or NANOG and SOX2 (C) expression in the indicated cell lines and timings of dox treatment. (C, E) Quantification of the results in B,D. Error bars indicate standard error of the mean (SEM). (F) Schematic of the experimental conditions for WNT induction in neural differentiation. (G, I) Immunostaining of OCT4 and NANOG (G) or SOX2 and BRA (I) after 6 days of dox induction during neural differentiation. (H, J) Quantification of the results in G and I, respectively. (K, L)

Bulk RNA-seq data from neural ectoderm differentiation assay on pluripotency marker expression at 2 (K) or 6 (L) days.

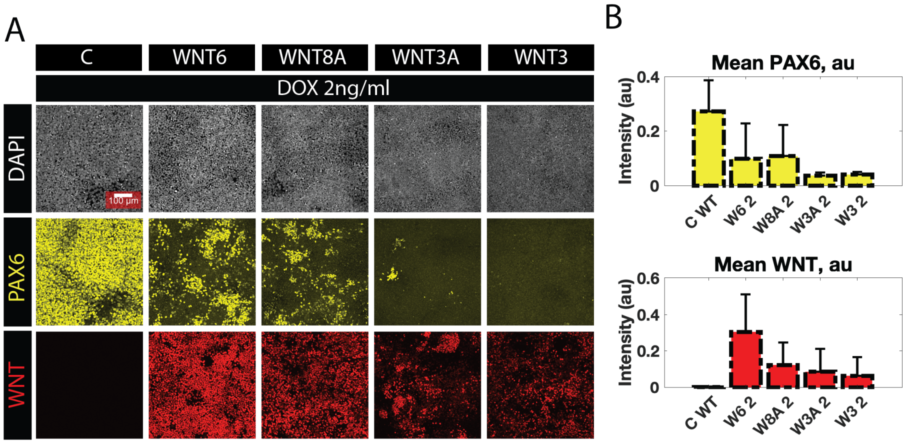


**Figure S4. PAX6 is downregulated by the four WNT ligands with different strengths.**
(A). Immunostaining for PAX6 in the indicated cell lines after 6 days of differentiation with DOX
(B). Quantification of the images in A. Error bars are the SEM values calculated from all the images per condition.

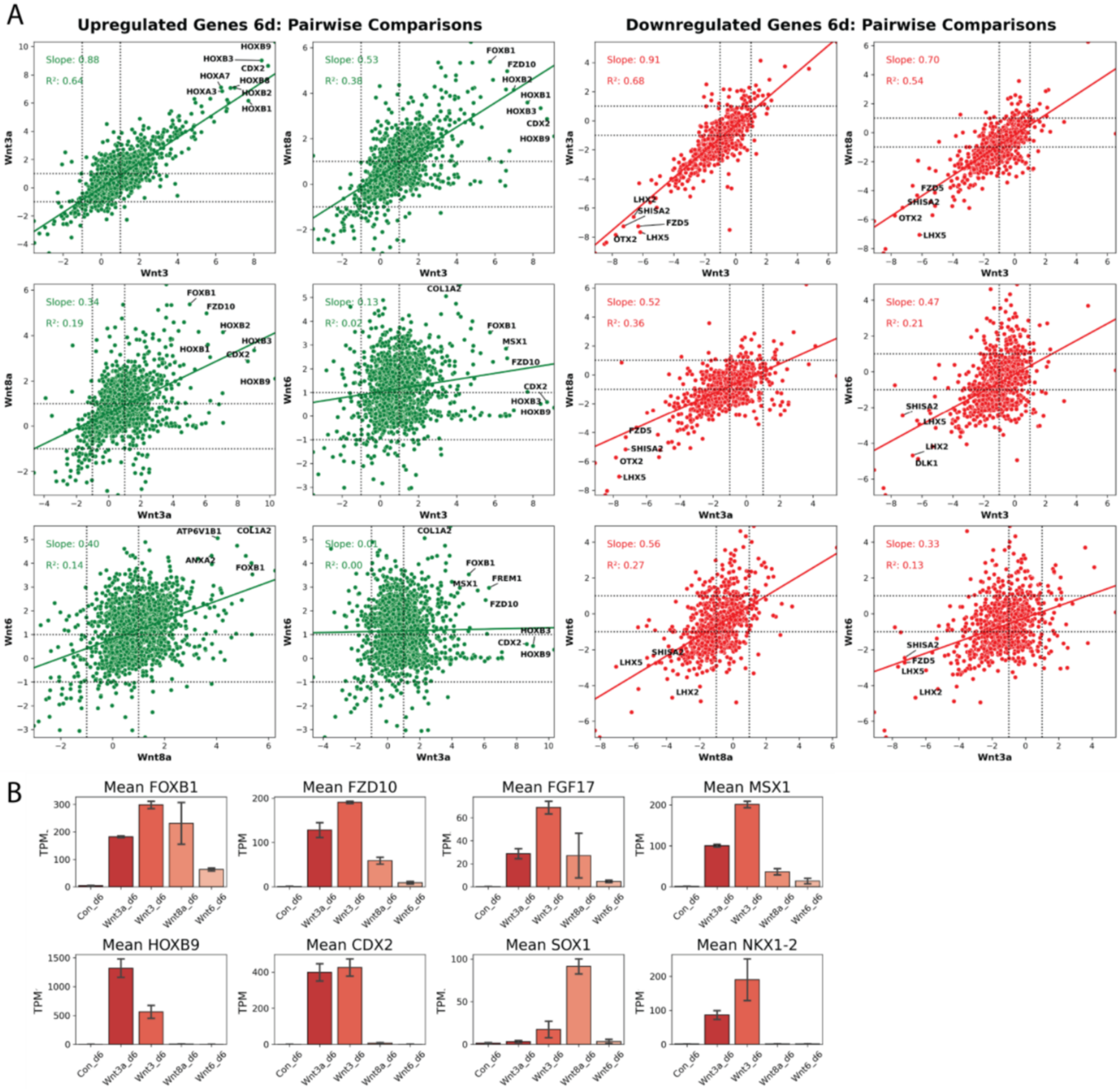


**Figure S5: Pairwise comparisons of gene expression induced by WNT3, WNT3A, WNT8A and WNT6.**

(A) Co-expression scatterplots with linear fits for upregulated (left) and downregulated (right) genes on day 6 under neural ectoderm differentiation. Genes that were upregulated or downregulated by at least 2-fold by one of the WNT ligands were included in plots on the left and right, respectively. (B) Expression of selected WNT target genes on day 6.

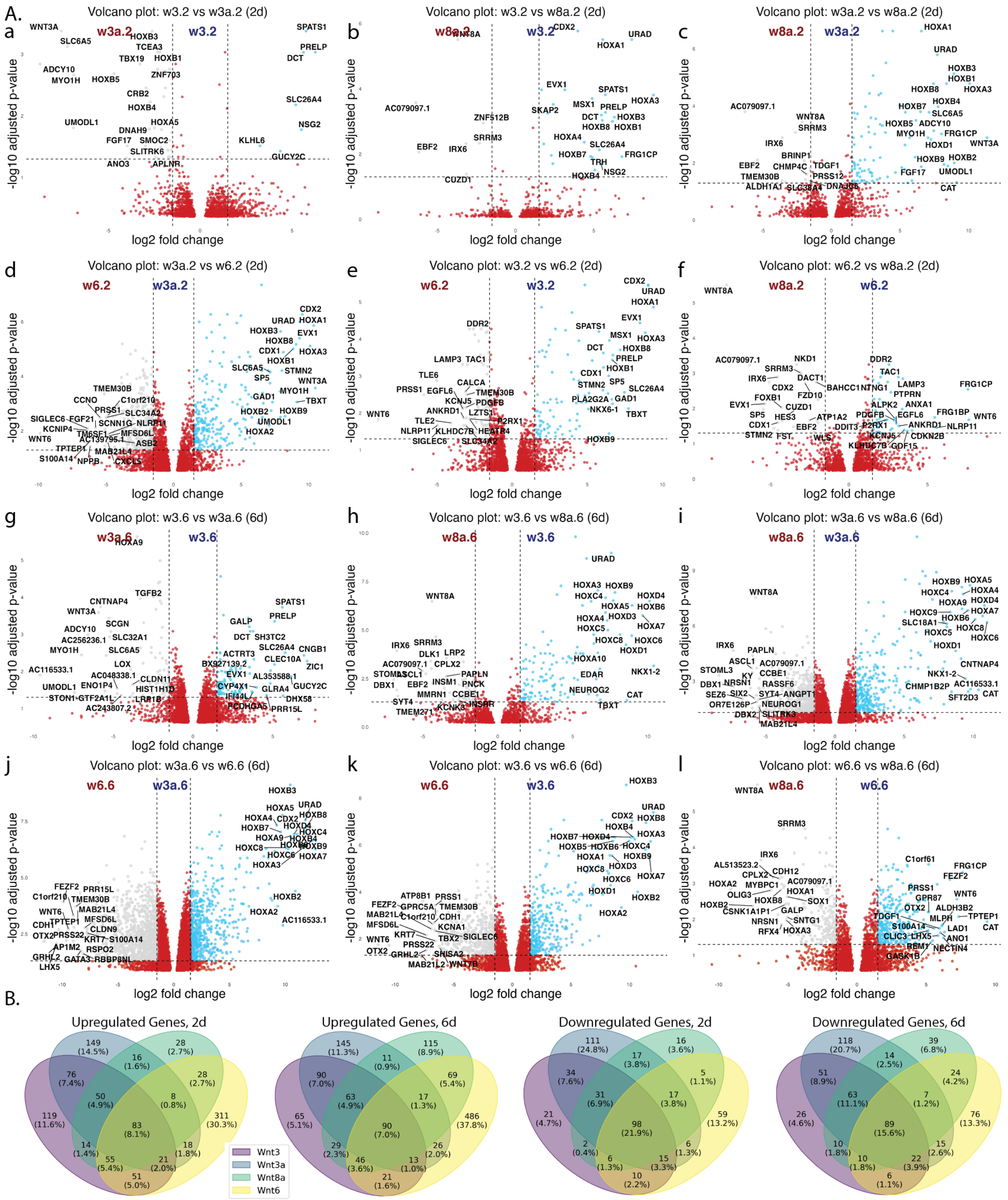


**Sup Figure 6: Differential expression analysis of WNT gene pairs on both day 2 and day 6**
(A). Pairwise Volcano plots show the differentially expressed genes between each pair of WNT ligands. (B). Venn diagrams of the number of common upregulated and downregulated genes across the four Wnt ligands. In all plots a cutoff of 2 fold change for differentiation expression was used.

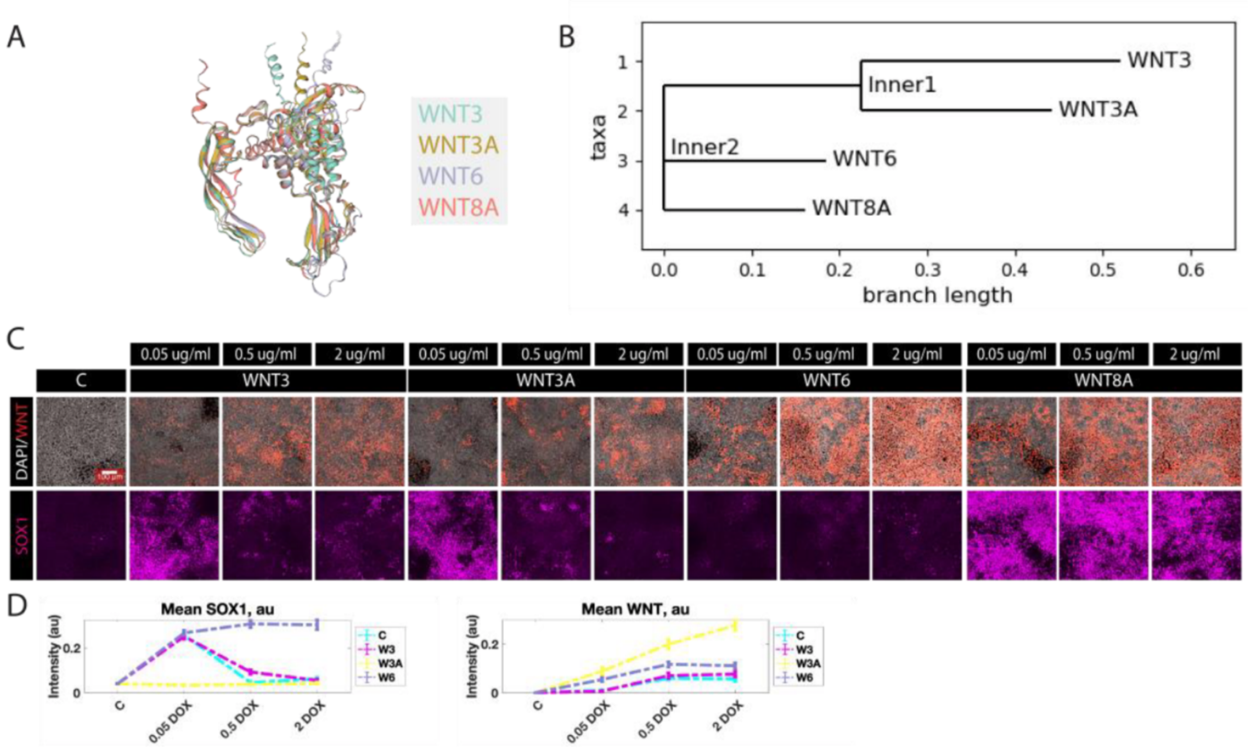


**Figure S7: WNT3 and WNT3A resemble each other structurally and functionally and suppress SOX1 at higher doses.**

(A). Structural superposition of WNT3, WNT3A, WNT6, and WNT8A highlighting key regions of variation. The index finger, thumb domain, and terminal regions show the highest structural divergence.

(B). Phylogenetic tree based on nucleotide sequence similarity of the four WNT genes, showing that WNT3 and WNT3A are the most closely related.

(C). Immunostaining for SOX1 expression in the indicated cell lines and doses of doxycycline during neural ectoderm differentiation.

(D). Dose–response curves showing SOX1 expression as a function of doxycycline concentration for each WNT ligand. Error bars indicate standard error of the mean between replicates (SEM).

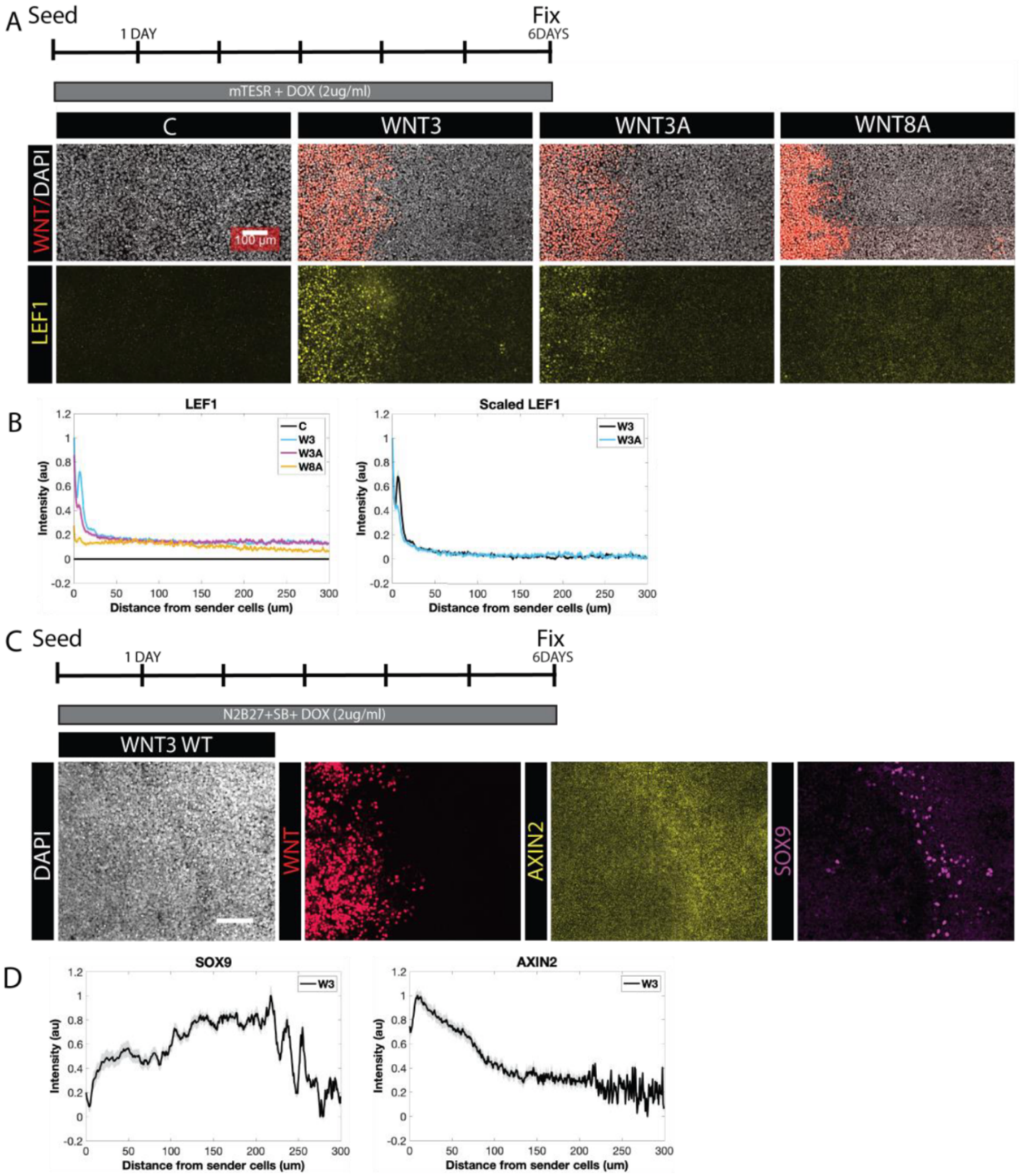


**Figure S8: Spatial Range of WNT ligand activity in epiblast or neuroectoderm differentiation assays.**

(A). Immunostaining of LEF1 expression induced in wildtype receiver cells by the indicated sender cells. Experiment performed in mTeSR1 pluripotency media.

(B) Quantification of LEF1 as a function of distance from the sender cells for the indicated cell lines. (C) Immunostaining of AXIN2 (a WNT inhibitor) and SOX9 expression domains induced WNT3 sender cells. (D) Quantification of the expression of AXIN2 and SOX9 as a function of the distance from W3 sender cells.

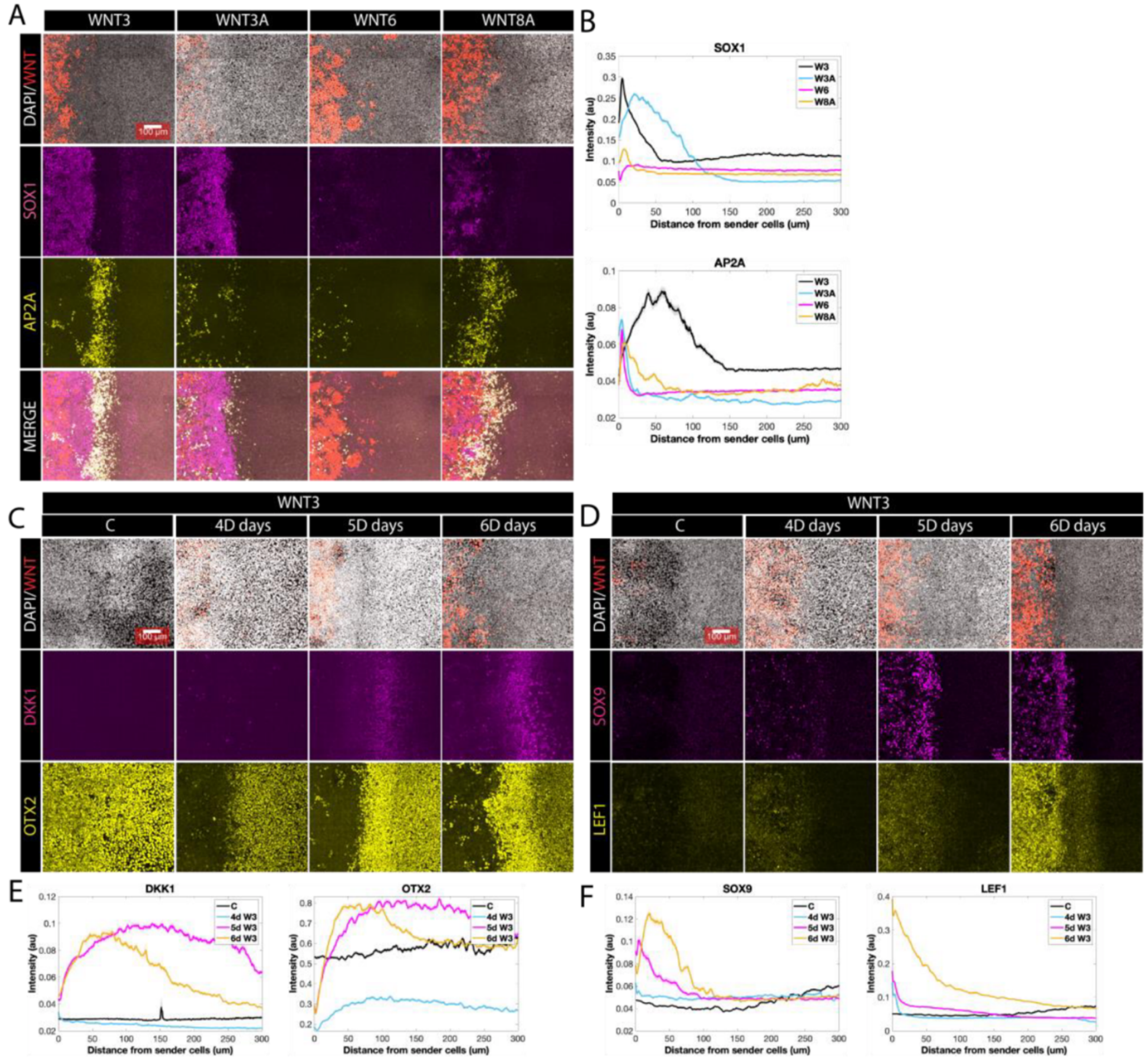


**Figure S9: Spatial patterns induced by localized WNT expression.**

(A) Immunostaining of SOX1 and AP2A in a 6-day juxtaposition assay under neural induction conditions for the indicated cell lines. Scale bar 100 um.

(B) Quantification of the expression of SOX1 and AP2A as a function of the distance from the sender cells.

(C, D) Immunostaining for the indicated markers on days 4, 5, 6 of the juxtaposition assay with the WNT3 overexpression cell lines. (E,F) Quantification of the indicated markers as a function of the distance from the sender cells.

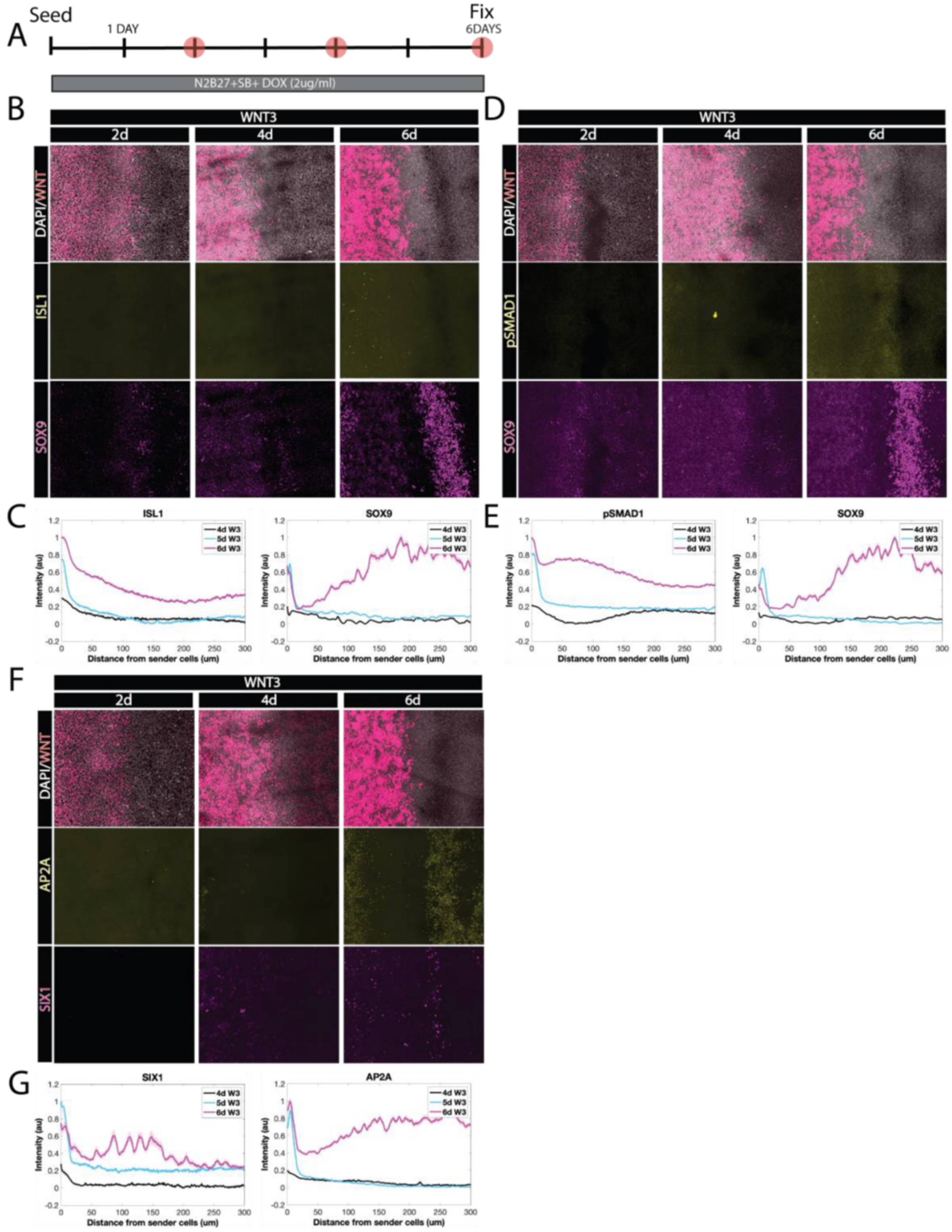


**Figure S10: Induction of spatial gradients in time by WNT3.**

(A). Schematic of 6-day juxtaposition assay under neural conditions.

(B, D, F). Immunostaining of the WNT3 juxtaposition experiments for the indicated markers and times. Scale bar 100 um.

(C, E, G). Quantification of the indicated markers as a function of the distance from the sender cells.
